# Predictive modeling of microbial data with interaction effects

**DOI:** 10.1101/2024.04.29.591596

**Authors:** Mara Stadler, Roberto Olayo-Alarcon, Jacob Bien, Christian L. Müller

**Affiliations:** Institute of Computational Biology, Helmholtz Zentrum München, 85764 Neuherberg, Germany; Department of Statistics, Ludwig Maximilians University Munich, 80539 Munich, Germany; Department of Data Sciences and Operations, University of Southern California, Los Angeles, CA, USA; Center for Computational Mathematics, Flatiron Institute, New York, NY 10010, USA

## Abstract

Microbial interactions are of fundamental importance for the functioning and the maintenance of microbial communities. Deciphering these interactions from (time-series) observational data or controlled lab experiments remains a formidable challenge due to their context-dependent nature, such as, e.g., (a)biotic factors, host characteristics, and overall community composition. Complementary to the classical ecological view, recent research advocates an empirical “community-function landscape” framework where an outcome of interest, e.g., a community function, is learned via statistical regression models that include pairwise or higher-order *statistical* species interaction effects. Here, we adopt the latter viewpoint and present penalized quadratic interaction models that can accommodate all common microbial data types, including microbial presence-absence data, relative (or compositional) abundance data from microbiome surveys, and quantitative (absolute abundance) microbiome data. We propose novel interaction models for compositional data and bring modern statistical techniques such as hierarchical interaction constraints and stability-based model selection to the microbial realm. To illustrate our framework’s versatility, we consider prediction tasks across a wide range of microbial datasets and ecosystems, including butyrate production in model communities in designed experiments and environmental covariate prediction from marine microbiome data. We show improved predictive performance of these interaction models and assess their limits in the presence of extreme data sparsity. On a large-scale gut microbiome cohort data, we identify interaction models that can accurately predict the abundance of antimicrobial resistance genes, enabling novel biological hypotheses about microbial community composition and antimicrobial resistance.

**Author Summary:** Microbes live in complex communities where interactions between species shape their function and stability. Understanding these interactions is crucial for predicting how microbial communities respond to environmental changes, medical treatments, or shifts in their host organisms. However, identifying these relationships is challenging because they depend on many factors, including the surrounding environment and community composition. In this study, we introduce a new statistical modeling approach to uncover microbial interactions from different types of data, including presence-absence patterns, relative abundance from microbiome surveys, and absolute abundance measurements. Our method builds on modern statistical techniques to improve accuracy and reliability, even when data are sparse or noisy. We demonstrate the power of our approach by applying it to diverse microbial datasets, from marine ecosystems to gut microbiomes. In one case, we successfully predicted antimicrobial resistance gene abundance based on microbial interactions, opening new avenues for understanding how resistance spreads in microbial communities. By advancing statistical tools for microbiome research, our work provides a new way to explore the hidden relationships between microbes, with potential applications in medicine, environmental science, and biotechnology.

## Introduction

A fundamental objective in microbial ecology is to elucidate how species compositions and species-species interactions are related to the maintenance and functioning of a microbial community [1]. Interactions between microbial species come in many forms, including cross-feeding interactions through metabolite exchange, bacteriocin-induced growth-inhibitory interactions, and exchange of genetic material for genotype selection [2, 3]. Conceptually, microbial interactions can be described in terms of their net positive, negative, or neutral effect on their interaction partner, resulting in broad categories such as mutualistic, commensal, or competitive interactions [4, 5, 6, 2]. Experimentally identifying and verifying such interactions within natural communities has remained a difficult task, owing to the sheer complexity of microbial ecosystems and limited technical capabilities to dissect such communities.

With the emergence of large-scale microbial survey data, computational approaches have become popular that use statistical regression and correlation methods to estimate sparse species-species association and co-occurrence networks from microbiome abundances [5, 7, 8, 9, 10, 11]. While these networks do *not* reflect true ecological relationships [12], they can provide valuable insights into the global structure of microbial communities across ecosystems [13, 14]. None of these methods, however, allow to relate species-species associations or “statistical interactions” to a community functional outcome of interest or to concomitant environmental or host-related covariates. Furthermore, most network approaches deliver context-independent (or averaged) pairwise associations, thus potentially missing species-species relationships that are relevant for a specific community function.

To enable context-specific microbial community modeling, the concept of *community-function landscapes* [15, 16] has been put forward as a promising empirical model for capturing how changes in microbial community composition affect collective function. The community-function landscape is essentially estimated from microbial data via statistical regression models that include pairwise or higher-order interaction terms. Here, we follow and extend this framework by using the generic quadratic interaction regression model as a starting point. We adapt this model to accommodate all common microbial abundance data modalities (see Fig. 1 for an illustration), such as, e.g., data from designed *in-vitro* experimental studies on model communities where microbes are given in presence-absence (binary) or absolute (count or continuous) abundance form [17, 18]. Importantly, our framework also extends to microbiome survey data where microbial compositions are measured by amplicon sequencing. These techniques provide relative abundance (or compositional) data [19] in form of Operational Taxonomic Units (OTUs) or Amplicon Sequencing Variants (ASVs) [20], or, when combined with absolute cell count measurements, (biased) quantitative microbial abundance information [21, 22, 23, 24].

**Fig 1.**
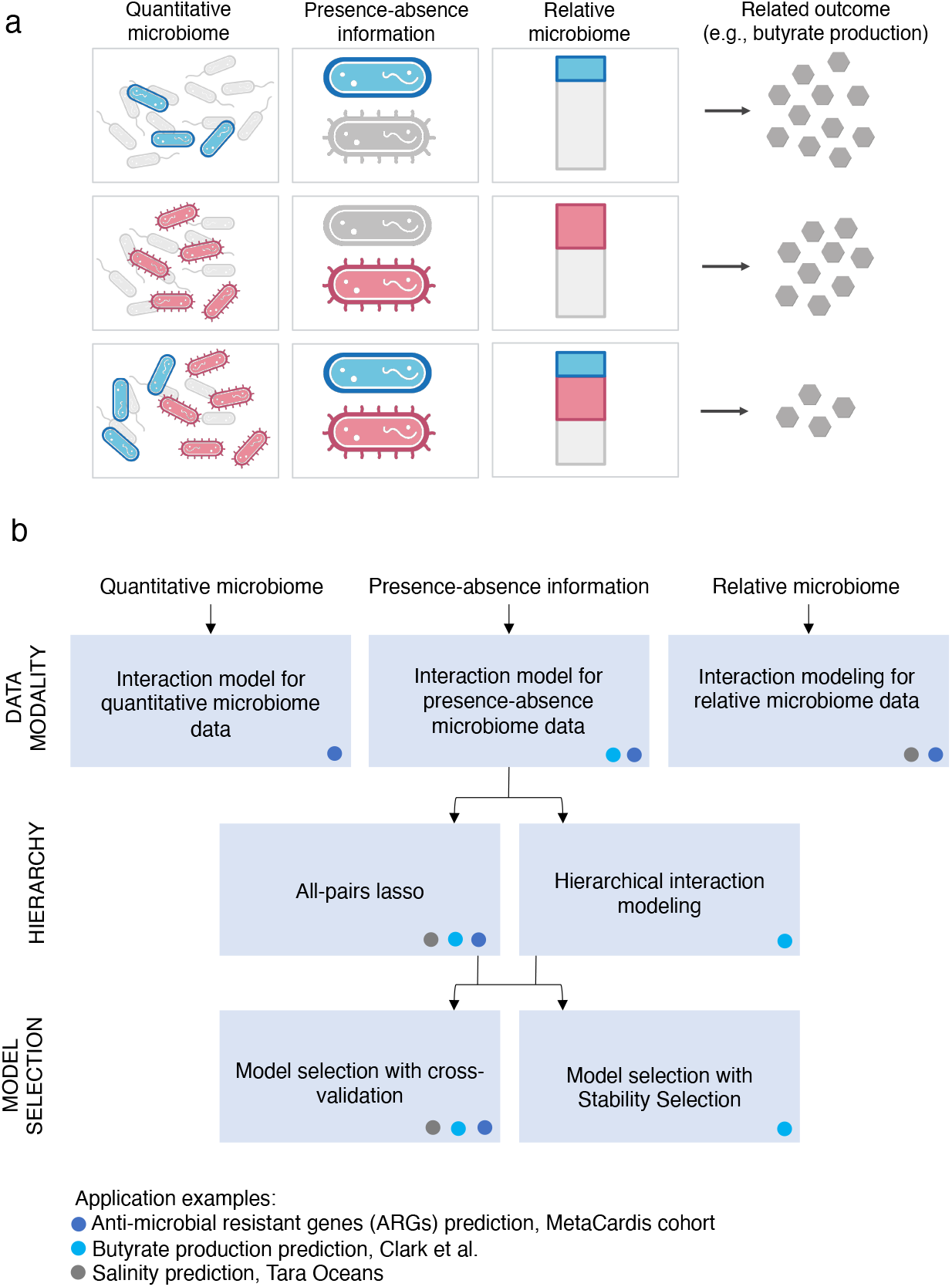
**a**. Illustration of the three most common data modalities in microbial data analysis and their combinatorial behavior with respect to an outcome, e.g., a community function: (i) quantitative microbiome data, representing absolute counts; (ii) presence-absence information of microbial species; and (iii) relative abundance (or “compositional”) data. Each row illustrates a simplified scenario. In the first scenario, blue microbes are present while red are absent, resulting in a large outcome (e.g., increased butyrate production). In the second scenario, red microbes are present while blue are absent, leading also to a large outcome. In the third scenario, both blue and red groups of microbes are present, yet only minimal amounts of the outcome are produced, indicating a potential interaction effect between the two species (created with BioRender.com). **b**. Illustration of the workflow modules (combinations of data and modeling options) presented in this study. Three real-world datasets (marked by the dots) are used to exemplify the respective modules.

Our framework unifies and generalizes several seemingly disjoint approaches in the microbial ecology and microbiome data science literature. On the one hand, it includes the community-function landscape view from microbial ecology where presence-absence and absolute abundance data from designed experiments on small microbial communities are used to predict community functions, such as, e.g., butyrate production, [18, 15, 16, 25] or overall host fitness [26]. Our framework is readily available for such studies and gives statistical guidelines how to choose model complexity, how to include additional constraints, and how to analyze higher dimensional datasets. On the other hand, our framework extends the linear regression model for compositional data, the so-called linear *log-contrast* model [27], to include statistical species-species combinations. Specifically, starting with Aitchison’s low-dimensional quadratic model [27], we introduce sparse quadratic interaction models that are applicable to large-scale relative abundance data derived from amplicon sequencing. To deal with the high dimensionality (where typically the number of taxa *p* and their pairwise interactions is larger than number of samples *n*), we follow the idea of penalized and structured log-contrast regression models [28, 29, 30, 31, 32, 33] and employ *ℓ*_1_ penalization on the interaction terms.

To deliver stable and interpretable interaction models [34, 35], we incorporate two key advancements from the high-dimensional statistics literature: (i) hierarchical interaction modeling [36, 37, 38] and (ii) stability-based model selection [39, 40]. The hierarchy assumption enforces constraints on interaction features, requiring them to be only included in the model if both features (strong hierarchy) or at least one feature (weak hierarchy) are already present as main effects. Stability-based model selection ensures that interactions are only included if they can be consistently and reproducibly identified across different subsets of the data, typically reducing the number of downstream testable hypotheses compared to standard cross-validation approaches.

We demonstrate the versatility of our framework by analyzing datasets that encompass all three data modalities across various ecosystems, including synthetic microbial communities, human gut microbiomes, and marine microbial ecosystems. Figure 1b presents an overview of the datasets used for the different workflow modules, i.e., combinations of regression models, model constraints, and model selection strategies. In the remainder of the paper, we introduce the statistical modeling strategies first (see **Methods**), followed by describing and discussing concrete microbial prediction tasks (see **Results**).

On quantitative human gut microbiome data from the Metacardis study [41] (Fig. 1b, dark blue dot), we show that the number of antimicrobial resistance genes (ARGs) can be well predicted by sparse interaction models on family-aggregated microbiome data, a considerable improvement over enterotype-based models [41].

On the Clark et al. [18] synthetic community dataset containing species presence-absence information and butyrate as community function, we identify the inhibitory role of *D. piger* on the butyrate producer *A. caccae* as the only stable interaction effect, considerably improving the interpretablity of prior community-function landscape models [18, 15].

For the *sparse quadratic log-contrast model* tailored for relative microbiome data, we (i) provide semi-synthetic data simulations to demonstrate the benefits and limitations of interaction effect inclusion and (ii) re-analyze Tara ocean data [42], achieving superior predictive performance on environmental parameters [33].

We conclude the paper by providing a comparative analysis of the interaction models across the three data modalities using the Metacardis ARG prediction task. We show that prediction quality decreases when using relative abundance or presence-absence data only and illustrate commonalities and differences between the learned models. The latter analysis gives guidance for the practitioner regarding the interpretability of quadratic interaction models. Our framework for microbial interaction modeling is available as reproducible R code at https://github.com/marastadler/Microbial-Interactions.

## Methods

### Interaction modeling strategy

Given the abundance information of *p* microbial taxa *X* = (*X*_1_, …, *X*_*p*_) across *n* samples, i.e., *X* _*j*_ ∈ ℝ ^*n*^, the baseline model for uncovering (joint) additive effects of the microbial taxa on an outcome *y* ∈ ℝ ^*n*^ (e.g., butyrate production), is the linear model

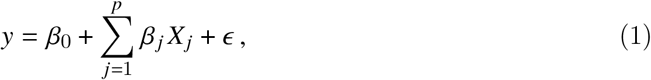

where *Β*_0_ ∈ ℝ is the intercept term, *Β* _*j*_ ∈ ℝ is the effect of taxon *j* on *y*, and *ϵ* models the technical and biological noise term. In many prediction tasks, relying on a linear (main effect) model alone is insufficient to accurately capture the community function or outcome of interest of the microbial community data. A common approach to introduce a more intricate yet interpretable model is the inclusion of quadratic terms. Here, we extend the baseline model by introducing a generic quadratic interaction model, incorporating all pairwise interactions between microbial taxa, namely

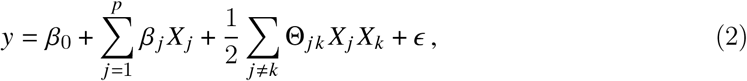

where Θ = Θ^*T*^ ∈ ℝ^*p*×*p*^ is a symmetric matrix of pairwise interactions. We assume the diagonal elements Θ_*jj*_ = 0 in this model formulation, though the general principles still apply if the constraint is removed. We next instantiate the interaction model in Eq. (2) to accommodate distinct data types and denote the microbial abundance information by ***A*** for count information (absolute or relative) and ***B*** for presence-absence information (see Fig. 1a).

#### Interaction model for quantitative microbiome data

Whenever microbial abundance information is given as absolute counts, the model is equal to the generic model 2 and does not require any transformation of the input data or any constraints on the model coefficients. Throughout this work, we denote the absolute count input data by 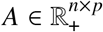.Assuming that *y* depends on the quantitative taxon abundances, the quadratic interaction model is given by

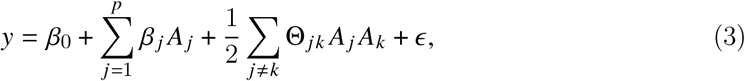

where the model parameters follow the description provided in Eq. (2).

#### Interaction model for presence-absence microbial data

If the microbial abundance information is represented as presence-absence data, we denote the resulting binary matrix as *B* ∈ {0, 1}^*n*×*p*^, where 1 indicates the presence of a microbial taxon, and 0 indicates its absence. One common alternative encoding is *B* ∈ {−1, 1}^*n*×*p*^, where the absence is encoded as -1. The choice of encoding does not affect the ability of the model to fit data but changes the *interpretation* of the coefficients. Assuming that *y* depends on the presence-absence information of microbial taxa, the quadratic interaction model is given by

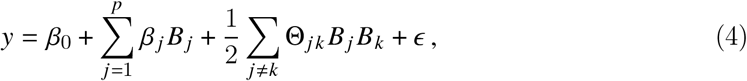

where the model parameters follow the description provided in Eq. (2). For *B* ∈ {0, 1}^*n*×*p*^, *Β*_0_ is the baseline effect when all features, i.e., microbial taxa, are absent, and *Β* _*j*_ for *j* = 1, …, *p* represents the effect of the presence of *B* _*j*_ when all other taxa are absent. The interaction term Θ_*jk*_ accounts for the additional effect when both features *B* _*j*_ and *B*_*k*_ are present. For *B* ∈ {−1, 1}^*n*×*p*^, *β*_0_ signifies the overall mean (assuming a completely balanced design). For more details on the interpretation and the linear transformations of model coefficients between these two encodings, see the Supplementary information. When describing *y* as a community-function, fitness, or phenotypic landscape, the different encodings in the interaction model are often associated with Fourier and Taylor expansions. The estimated parameters are then used for the construction of landscape descriptors, such as, e.g., “ruggedness” (see [15, 43, 44] for further details on such landscape analysis).

#### Interaction modeling for microbiome relative abundance data

One of the most abundant microbial data sources is amplicon sequencing where the derived taxa counts carry only relative abundance information. Typically, each sample, i.e., each row of *A*, is represented as a compositional vector whose entries sum up to a constant [7, 8, 19]. This implies that all features are necessarily statistically dependent, thus requiring adequate data transformations for linear and quadratic interaction modeling, respectively. One popular way, put forward in compositional data analysis [27], are so-called log-ratio transformations. The additive log-ratio transformation (alr) requires a “reference feature”, e.g., the *p*th feature, and builds log-ratios with respect to that reference (see [45] for a detailed discussion of the alr transform). The transformed count vector is given by *C* _*j*_ = log(*A* _*j*_ /*A*_*p*_), *j* = 1, …, *p* − 1. The alr transformation allows principled linear modeling of an outcome *y* using the (*p* − 1)- dimensional log-ratios as features via:

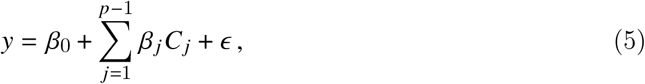

where the coefficient *β* _*j*_ quantifies how the outcome *y* is related to the log-differences of the *j* th feature with respect to the chosen reference. A natural extension for quadratic interaction modeling is thus the so-called *alr transformed quadratic model*, given by

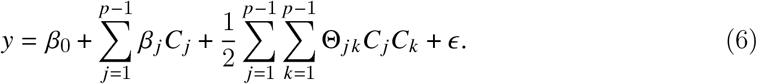

For *p* := *p* − 1 this formulation is equivalent to the generic model in Eq. (2) (with Θ not being symmetric). While this model formulation allows for the interpretation of the effects with respect to the *p*th reference feature, a more convenient reference-free symmetric expression of the linear alr transformed model in Eq. (5) can be derived by reformulating the equation as a *p*-dimensional problem with a zero-sum constraint, given by

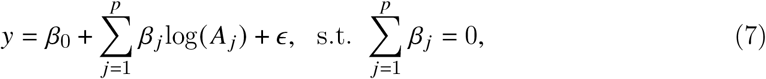

where the main (log) effect coefficients *β* _*j*_, *j* = 1, …, *p* sum up to zero. As illustrated in [27], this so-called linear *log-contrast* model can be extended to the quadratic log-contrast model as follows:

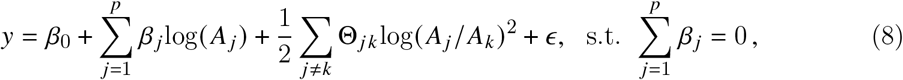

where the main (log) effect coefficients *β* _*j*_, *j* = 1, …, *p* sum up to zero, with *β* ∈ ℝ^*p*^, and the interaction effect coefficients Θ_*jk*_ correspond to the quadratic (log-ratio) interaction effect of *A* _*j*_ and *A*_*k*_, with Θ = Θ^*T*^ ∈ ℝ^*p*×*p*^. We denote this model as the *constrained quadratic log-contrast model*. In the Supplementary information we show how to formulate the alr transformed model as constrained quadratic log-contrast model. For completeness, we also demonstrate in the Supplementary information how the interaction model for all pairs of log-ratios [31] is defined for compositional data.

### Penalized model estimation

Microbial datasets typically include a large number of features *p* and interactions between features *p* (*p* − 1)/2 compared to the number of observations *n*. Even in scenarios where *n > p* (*p* − 1)/2, a parsimonious model is often more appropriate, enabling the selection of only a few features and interactions that are most relevant for the outcome.

To facilitate penalized model estimation, we employ regularized maximum-likelihood estimation incorporating *ℓ*_1_-norm (lasso) penalization [46] for both linear and interaction coefficients. We introduce a generic optimization problem, consisting of an objective function *ρ*_*λ*_ (*l, β*_0_, *β*, Θ) and a (potential) constraint set on the model parameters *c*(*β*_0_, *β*, Θ) that facilitates parameter estimation for all (linear and interaction) models introduced before. The objective function takes the general form

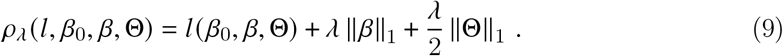

Here, *λ >* 0 serves as a tuning parameter, regulating the sparsity levels of the coefficients *β* and Θ, respectively. The loss function *l* (*β*_0_, *β*, Θ) is specific to each model. Consequently, the generic optimization problem is given by

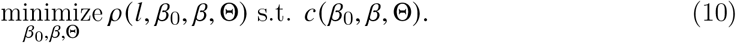

This optimization problem is now instantiated by specific loss functions and constraints.

#### Sparse quadratic interaction model for quantitative and presence-absence micro-biome data

The loss function *l* (*β*_0_, *β*, Θ) for the sparse quadratic interaction model, also known as all-pairs lasso, for the interaction models for absolute count data or presence-absence data, introduced in Eq. (3) and Eq. (4), is defined as

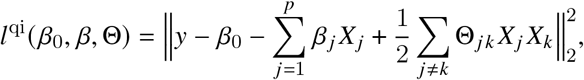

with 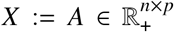 for absolute count data and *X* := *B* ∈ {0, 1}^*n*×*p*^ (or *B* ∈ {−1, 1}^*n*×*p*^) for presence-absence data. This model does not require further constraints on the model parameters, such that *c*(*β*_0_, *β*, Θ) = ∅. Consequently, the optimization problem is given by

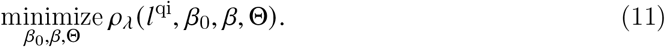

In the linear model case the loss function in the optimization problem reduces to 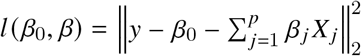

#### Sparse quadratic log-contrast model

While the linear log-contrast model has been extended to the high-dimensional setting via sparsity-inducing penalization [28, 29, 32, 33], the quadratic log-contrast model [27] has not yet been applied to any high-dimensional data. We thus define the loss function for the sparse quadratic log-contrast model (qlc) for compositional data, introduced in Eq. (8) as

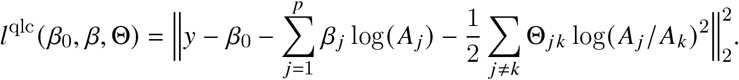

Since this model includes a zero-sum constraint on the main effect coefficients, the constraint set in Eq. (10) is given by

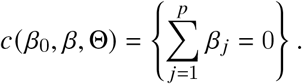

Thus, the optimization problem for the sparse quadratic log-contrast model is given by

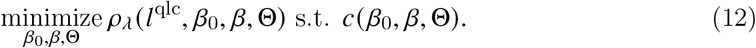

In the linear sparse log-contrast model (lc) defined in Eq. (7), the loss function reduces to 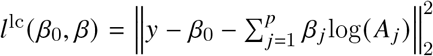 The corresponding optimization problems can be efficiently solved with the `c-lasso` solver [47] as integrated in the R package `trac` [33, 48]. In the linear log-contrast model, the main effect covariates log(*A* _*j*_) for *j* = 1, …, *p* do not require scaling since the model is equivariant under the zero-sum constraint (see [33] for an outline of this property). For the quadratic log-contrast model, however, we require proper scaling of the interaction features log(*A* _*j*_ /*A*_*k*_)^2^ to ensure a balanced effect of penalization with respect to the main effects. While different scaling procedures are conceivable, we propose the following centered log-ratio (clr) scaling. The clr divides each compositional part by the geometric mean of all parts, namely

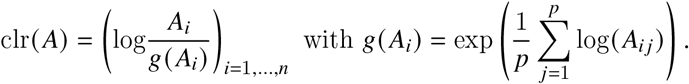

To achieve a balanced penalization effect, we apply scaling to align the *ℓ*_2_-norms of the interaction features with the average *ℓ*_2_-norm of the main effects after clr transformation. This approach ensures consistent penalization across both main and interaction terms, improving model interpretability and regularization.

Mathematically, this can be expressed as follows. We denote each column of the interaction feature matrix as 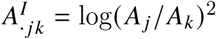, with *A*^*I*^ ∈ ℝ^*n*×*p*(*p*−1)/2^, and its scaled version is given by

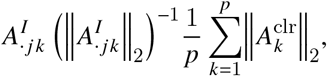

where *A*^clr^ = clr(*A*) ∈ ℝ^*n*×*p*^ is the clr transformed main effects matrix *A* and 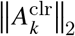 is the *ℓ*_2_-norm of the *k*-th column of *A*^clr^. For completeness, we detail the optimization problems for the sparse additive log-ratio (alr) transformed quadratic model and the sparse quadratic log-ratio model in the Supplementary information.

### Modeling hierarchical interactions

While quadratic interaction models can generally enhance predictive performance compared to the linear counterparts, interpretability of the resulting models becomes more challenging. To strike a good balance between prediction quality and interpretability, we introduce the statistical concept of hierarchy in the context of quadratic models for microbial data. The concept of hierarchy permits the inclusion of an interaction Θ_*jk*_ in the model *only if* both associated main effects are also present in the model (strong hierarchy), or if at least one of the associated main effects is included (weak hierarchy) (see [36], and references therein, for further discussion). For many microbial consortia, it is reasonable to assume that bacterial species only show an empirical interaction effect if each of them has an independent effect on community function. For example, two species may independently contribute to butyrate production but compete for the same limited food source, thus resulting in a potentially negative interaction effect. This hierarchy principle can be implemented by imposing constraints on the interaction effects Θ_*j*_ ∈ ℝ^*p*^ for *j* = 1, …, *p* as follows:

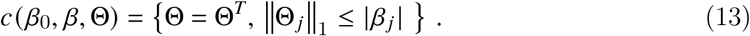

By eliminating the symmetry constraint on Θ, the resulting model relaxes to weak hierarchy on the interaction features. Moreover, this approach allows a strong interaction to “pull” itself into the model, ensuring that it cannot be missed, even if it violates the hierarchy assumption. While the constraint in Eq. (13) results in a non-convex optimization problem, we follow [36] who proposed a convex relaxation of the problem and provided an efficient implementation in the corresponding R package hierNet [49] (v1.9). The hierarchical constraint can be imposed within the generic optimization problem described in Eq. (10) and can be readily included for (i) quantitative microbiome data (ii) or presence-absence microbial data in Eq. (11), and (iii) relative abundance data in the alr-transformed interaction model in Eq. (S.3).

### Model selection

An essential challenge in high-dimensional penalized regression is the selection of the regularization parameter *λ >* 0. This parameter balances the sparsity of model coefficients with out-of-sample predictive performance [50, 51]. Standard methods for main effects and interaction models often involve Information Criteria like the Akaike (AIC) and the Bayesian Information Criterion (BIC) [52]. Here, we revisit standard cross-validation techniques [36] and introduce stability-based model selection for quadratic interaction models.

#### Cross-validation

The principle idea of cross-validation is to split the training data set of size *N* (typically 50 − 80% of the available sample size *n*) into multiple folds where each fold is used both for learning the model parameters and as in-sample validation set. In leave-one-out cross-validation (LOOCV), the model is trained *N* times, with each training using all data points except one, and the left out data point is used for model validation. Skwara et al. [15] used this approach to learn community-function landscapes. *K*-fold cross-validation (CV) divides the training dataset into *K* subsets (or folds), training the model on *K* − 1 of these folds and using the remaining fold for validation. This process is repeated *K* times such that each fold serves once as the validation set. This approach helps find the *λ* that minimizes the prediction (or classification) error, effectively balancing bias and variance. In regression, the chosen *λ*_MSE_ minimizes the average mean-squared error

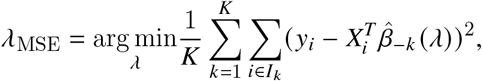

where *I*_*k*_ denotes the subset of observations without fold *k*. An alternative sparsifying heuristic is to choose the largest *λ* that is within the one standard error (1se) band of the average mean-squared error at *λ*_MSE_, denoted by *λ*_1se_. To reduce the dependency of model selection on specific data splits, the cross-validation procedure can be repeated multiple times, enabling the construction of empirical coefficient distributions (as shown in **Results**).

#### Stability selection

Cross-validation is known to overselect coefficients in sparse regression [52]. To guard against this false positive coefficient selection, stability selection [39] has been introduced as popular alternative. Stability selection has shown effectiveness across various scientific domains, ranging from network learning [53, 54] to data-driven partial differential equation identification [55, 56]. In linear regression, stability selection involves iteratively learning sparse regression models from subsamples of the data (e.g., *n*_*s*_ = ⌊*n*/2⌋), recording the frequency of selected predictors across models, and selecting the most frequent predictors for the final model. A variant of stability selection, complementary pairs stability selection (CPSS) [40], is particularly advantageous for handling unbalanced experimental designs, as it ensures that individual subsamples are independent of each other. CPSS draws *b* subsamples as complementary pairs {(*a*_2*l*−1_, *a*_2*l*_) : *l* = 1, …, *b*}, indexed by *a*_2*l*−1_ and *a*_2*l*_, where *l* denotes the pair number, with *a*_2*l*−1_ ∩ *a*_2*l*_ = ∅ from samples {1, …*n*} of size ⌊*n*/2⌋. For each subsample, a sparse model is learned using a variable selection procedure *S* which influences stability and complexity of the resulting model [39]. A popular selection procedure is to pick the *k* first predictors that enter the model along the regularization path (also referred to as *λ*-path). Applying a variable selection procedure *S* to each subsample allows defining a feature-specific selection probability 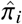 for *i* = 1, …, *p* + *p* (*p* − 1)/2 that is given by

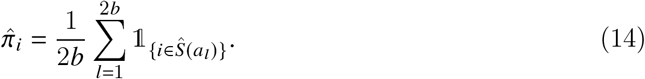

The final selection set, denoted as *Ŝ*^CPSS^, consists of predictor indices *i* for which the estimated selection probability 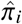 exceeds a predefined threshold π_thr_, that represents the minimum selection frequency required for a predictor to be included in the final set. We employ the stabs R package [57] (v0.6-4), which offers an efficient implementation of the CPSS procedure. This approach involves defining several hyperparameters, including the set of regularization parameters Λ, the threshold π_thr_ ∈ [0, 1], the number of initial predictors *k* entering the sparse model, and the number of complementary splits *b*. The CPSS procedure in [57] can be directly applied to linear models. Since CPSS does not make a distinction between main and interaction effects, we apply CPSS also to quadratic models with and without hierarchical constraints [35].

## Results

### Quantitative genus-level interaction models can predict antimicrobial resistance gene abundances

Large-scale metagenomics survey data enable not only the quantification of species compositions but also the abundances of genes and pathways associated with a particular function or metabolic potential [58]. Here, we investigate the question whether the number of antimicrobial resistance genes (ARGs) in a community, as identified from the respective community metagenomes, can be predicted from high-level taxonomic compositions. A good taxonomic-based statistical interaction model would allow to assess the antimicrobial potential of a microbial community when only quantitative amplicon data are available, [21]. Prior work has shown that there are significant associations between human gut enterotypes, i.e., distinct community types that are dominated by specific genera in the population [59], and the number of ARGs across countries [60]. Recent studies not only report significant associations between the number of ARGs and community diversity but also link specific species groups to ARG abundance and horizontal gene transfer [61].

We base our analysis on metagenomic Operational Taxonomic Units (mOTUs) abundances (*p*_s_ = 1938 species) from quantitative microbiome profiling for a subset of *n* = 690 individuals from the MetaCardis cohort for which the number of identified ARGs *y* ∈ ℝ^*n*^ are available [41]. We consider three statistical regression models: (i) enterotype-based regression where individuals are binary-encoded to one of the four enterotypes *Bacteroides 1* (high percentages of Bacteroides and Faecalibacterium), *Bacteroides 2* (high percentages of Bacteroides and low percentages of Faecalibacterium), *Prevotella* (high percentages of Prevotella), and *Ruminococcus* (low percentages of Bacteroides and Prevotella) (see also [62]), (ii) sparse linear regression (Eq. 1), and (iii) a (sparse) quadratic interaction model (Eq. 2). For the latter two models, we aggregate the *p*_s_ = 1938 species to genus level and collect the *p* = 30 most prevalent genera (out of 720) as potential predictors in *A* ∈ ℝ^690×30^. The enterotype-encoded model is solved without regularization, using a standard linear model. The linear model is solved using regularization and 5-fold CV with minimum MSE *λ* selection. Since the microbial counts in *A* are quantitative, we fit the sparse interaction model for absolute count data defined in Eq. (3) and Eq. (11), respectively, by using 5-fold CV with minimum MSE *λ* selection. All three models are trained and evaluated using the same 10 train-test splits, with 2/3 of the data allocated for training and 1/3 for testing.

We observe that enterotype-based predictions show remarkable predictability for ARG abundances given the simplicity of the model (median out of sample *R*^2^ 0.36) (see Fig. 2a,b). The model reveals that the number of ARGs are positively associated with the Bacteroides enterotypes and that both the *Prevotella* and *Ruminococcus* enterotypes are negatively associated, respectively (see also Fig. S1 for a visual representation). The genus-based linear model considerably improves predictability (median out of sample *R*^2^ 0.48, see Fig. 2b). Consistent with the enterotype-based model, we observe a strong positive effect on Bacteroides and a strong negative effect on Prevotella, respectively (see Fig. S2). The linear model, however, reveals a more fine-grained picture of the empirical dependencies with strong positive effects of Escherichia, and Parabacteroides, and negative effects of Faecalibacteria and Phascolarctobacteria on ARG abundance (see Fig. S2). Strikingly, the sparse quadratic interaction model achieves significantly higher out of sample *R*^2^ consistently above 0.5 (see Fig. 2c for a typical out-of-sample prediction). Figure 2d shows the nine stable linear effects and 14 stable empirical interactions among genera that have a non-zero median effect over 10 train test splits. The two strongest positive interactions are between Prevotella and Dorea, and Prevotella and Faecalibacteria, respectively. The latter relationship indicates an empirical “antagonistic” association between the two genera, given their individual negative effects. These findings suggest that presence *and* co-presence of certain bacterial genera in the gut microbiota can predict community-wide prevalence of ARGs.

**Fig 2.**
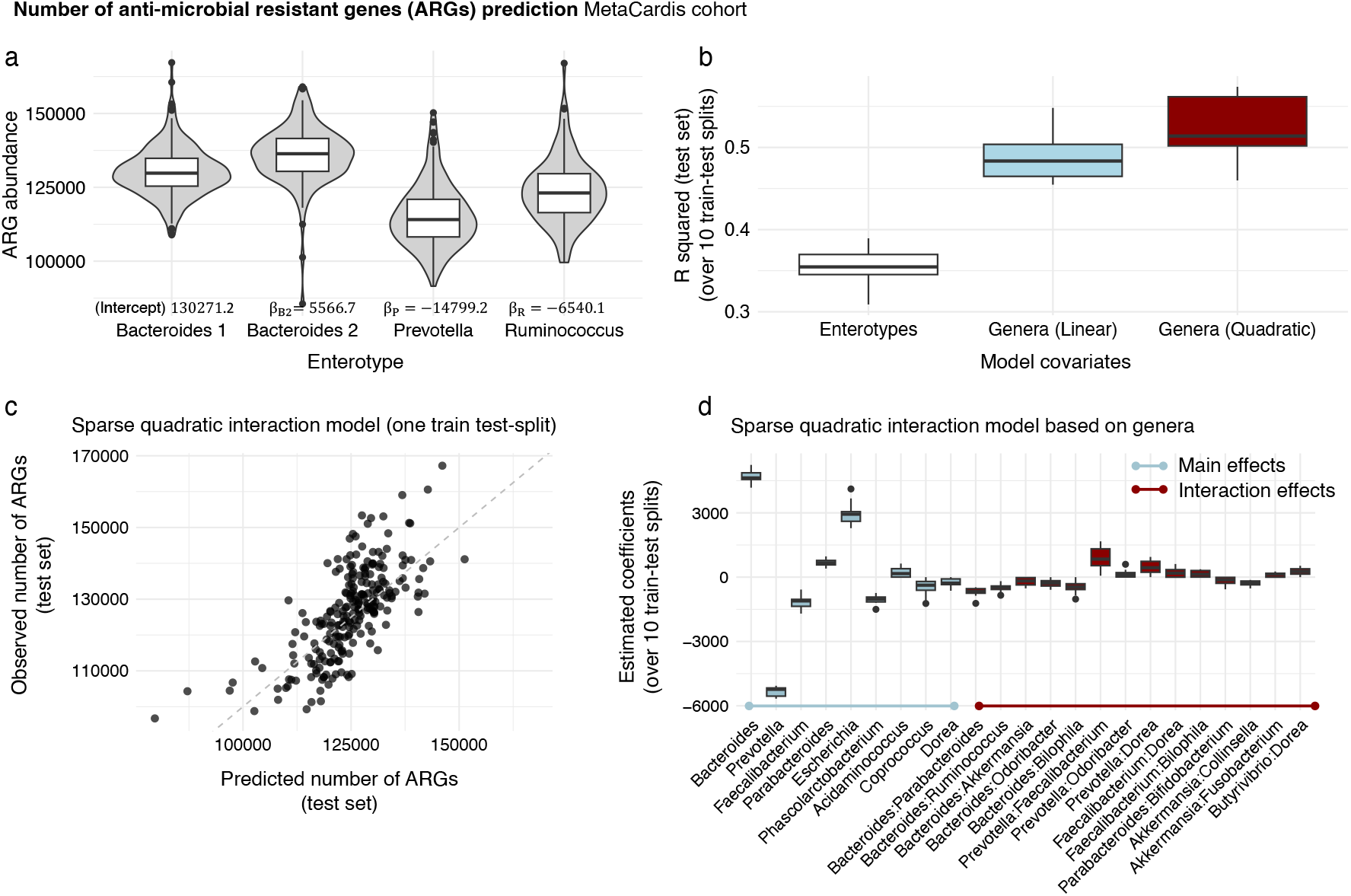
Prediction of antimicrobial resistance gene (ARG) abundances from enterotype and genus-level absolute abundance information from the MetaCardis cohort [41]: **a**. Distribution of ARG abundance per enterotype (Bacteroides 1, Bacteroides 2, Prevotella, and Ruminococcus). Significant coefficient estimates from the linear model are shown below the boxes. **b**. Distribution of test set *R*^2^ (10 splits) for ARG prediction based on enterotype or genera abundances (sparse linear and quadratic models, respectively). **c**. Observed vs. predicted number of ARGs on a representative test data set for the sparse quadratic interaction model. **d**. Distribution of estimated coefficients with non-zero median over 10 train test splits in the sparse quadratic interaction model 11.

### Hierarchy and stability-based model selection improve model interpretability for community-function landscapes

We next investigate the community-function landscape of an in-vitro bacterial community comprising *p* = 25 members with respect to the production of butyrate, a short-chain fatty acid beneficial to human health [18]. Butyrate-producers have the ability to ferment dietary fibers into butyrate, contributing to gut health, immune function, and energy metabolism [63]. Following [15], we use the published presence-absence data from the *n* = 1561 designed experiments by Clark et al. [18], denoted by *B* ∈ {0, 1}^*n*×*p*^, to fit sparse quadratic interaction models defined in Eq. (4) and Eq. (11) for predicting butyrate production *y* ∈ ℝ^*n*^.

Prior analysis of these data has shown that sparse linear, quadratic, and third-order interaction models with LOOCV tuning can increasingly well approximate the community-function landscape for butyrate production with in-sample and LOOCV *R*^2^ around 0.7 − 0.9 (see [15], Fig. 2a for reference). Here, we specifically investigate the effect of hierarchical constraints on quadratic interaction inclusion and the choice of the model selection on predictive performance, model complexity, and interpretability. As summarized in Fig. 1b, we used the all-pairs lasso formulation and hierarchical interaction modeling combined with 5-fold CV and CPSS for model selection. We used ten random train-test splits where, for each split, 2/3 of the data were used for training/model selection, and 1/3 for measuring out-of-sample test performance. Our analysis revealed that, similar to [15], the sparse quadratic interaction model without hierarchical constraints and 5-fold CV yield excellent predictive performance with a median out-of-sample *R*^2^ of 0.78. Including hierarchical constraints slightly reduced the performance in terms of median out-of-sample *R*^2^ to 0.72. Figure 3 illustrates the benefit of hierarchical modeling by greatly reducing model complexity. The sparse quadratic interaction models comprise, on average, 152.7 main and interaction effects whereas hierarchical models require, on average, 38.5 non-zero coefficients. Figure 3a shows the sorted median coefficients for both models, revealing a large number with small effect sizes for the all-pairs lasso solution (Fig. 3a, gray bars). Figure 3b focuses on the sorted 38 non-zero model coefficients in the hierarchical interaction model (green bars) and compares their effect sizes with those of the model without hierarchy. The hierarchical model posits a non-zero main effect on all 25 species in the community and includes 13 interaction effects. Both the hierarchical and standard quadratic interaction model identify *Anaerostipes caccae* (AC), a known acetate and lactate-utilizing bacterium [64], to have the strongest positive effect on butyrate production, followed by *Coprococcus comes* (CC) and *Eubacterium rectale* (ER). These results are consistent with the original analysis in Clark et al. [18]. However, only the hierarchical model correctly identifies *Desulfovibrio piger* (DP) to have a strong negative effect on butyrate production (second to last bar in Fig. 3b).

**Fig 3.**
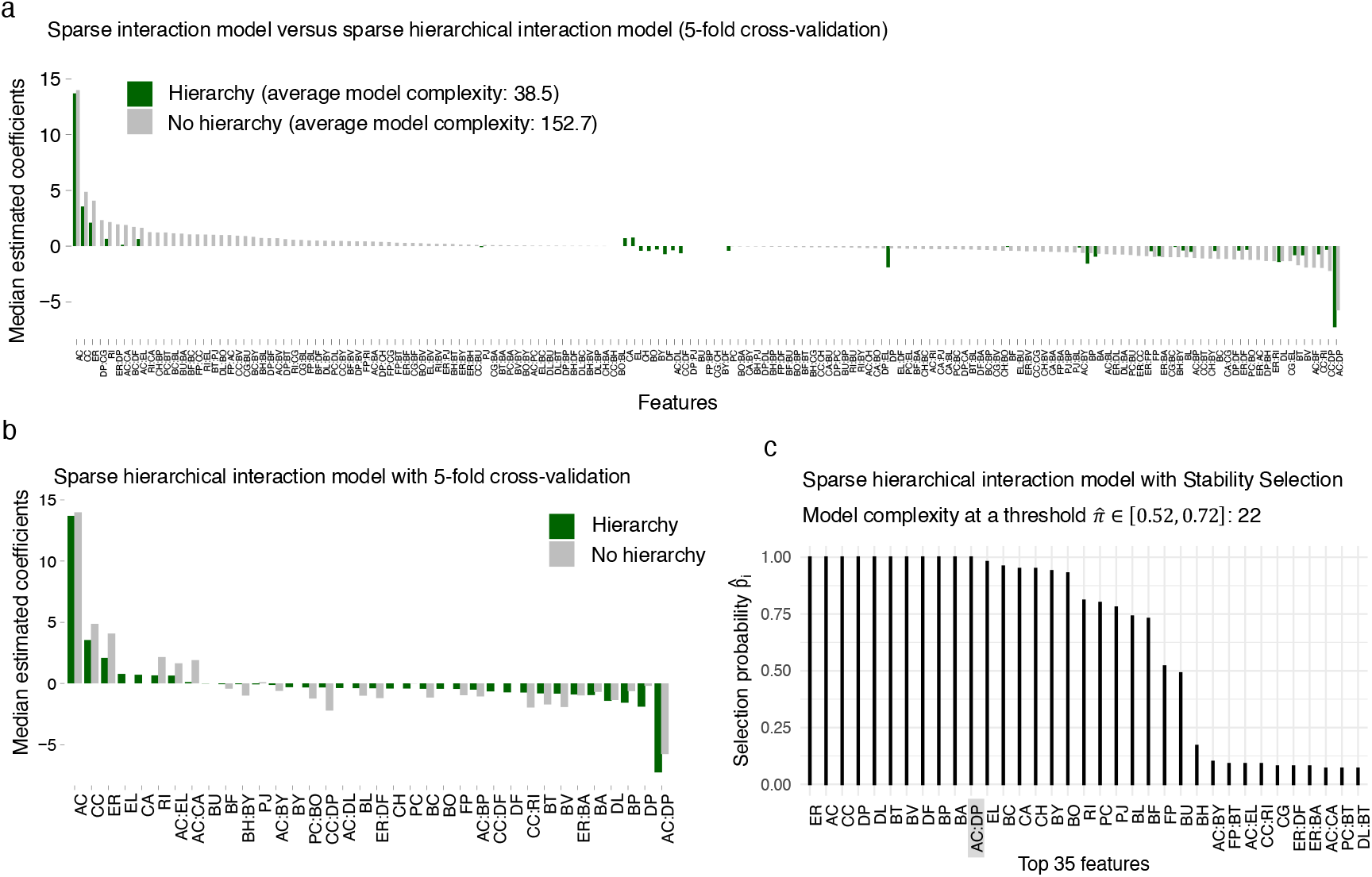
a. Comparison of median estimated coefficients (over 10 train test splits) between the sparse quadratic interaction model and the sparse hierarchical interaction model (Eq. (13)) with weak hierarchy. Only estimates that are non-zero in at least one model are shown. **b**. Median non-zero estimated coefficients for both models. Only coefficients that are non-zero in the hierarchical model are displayed. For direct comparison, the corresponding coefficients from the model without hierarchy are included. **c**. Top 35 selection probabilities from complementary pairs stability selection (CPSS) in the sparse hierarchical interaction model.

Among the thirteen interactions, jointly identified by both models, *Anaerostipes caccae* and *Desulfovibrio piger* (AC:DP) show the strongest negative interaction. The inhibiting effect of *D. piger* on *A. caccae*’s butyrate production, first observed in co-culture experiments [65], is likely related to hydrogen sulfide production by *D. piger* from lactate that, in turn, impacts *A. caccae*’s ability to produce butyrate [65]. Other interactions, including those between *A. caccae* and *Eggerthella lenta* (AC:EL), and *Collinsella aerofaciens* (AC:CA), help improve prediction, yet require further experiments regarding potential mechanism.

Moreover, when applying complementary pairs stability selection (CPSS) in place of cross-validation for hierarchical interaction modeling, model complexity is further reduced to 24 main and interaction effects, on average, without significant reduction in predictive performance. Figure 3c summarizes the stability profile in terms of CPSS selection probabilities 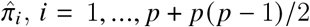.We observe that only 16 species main effects consistently appear in the models with selection probabilities 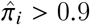 and only one interaction effect, the one between *A. caccae* and *D. piger*. Seven main effects have selection probabilities 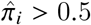 before the stability profile sharply drops off, with the remaining single species and interaction effects having negligible selection probabilities. These results demonstrate that state-of-the-art statistical modeling with hierarchy constraints and stability selection mechanisms allows a tremendous reduction in community-function models complexity without sacrificing predictive performance.

### Interaction modeling improves predictive performance for microbiome sequencing data

We next investigate the behavior and the performance of our novel quadratic interaction models for (sparse) high-dimensional relative abundance data from amplicon sequencing. Since the excess number of zero measurements as well as experimental noise in such data can hamper proper interaction modeling, we first perform a semi-synthetic simulation study that elucidates the conditions for which accurate estimation is feasible. This analysis is followed by an application study using the Tara global ocean microbiome data [42]. Rather than estimating a community-function landscape, we estimate a “community-environment” landscape with ocean salinity concentration as target environmental characteristic [42, 33].

#### Semi-synthetic simulation setup

To create realistic and flexible benchmark scenarios, we first resort to a semi-synthetic simulation framework where we leverage 16S rRNA amplicon sequencing data from the American Gut Project [66]. We use the pre-processed American Gut data from [33], comprising *n* = 6266 samples and *p* = 1387 OTUs with associated taxonomic information. The compositional count matrix 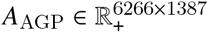 comprises taxa with a high variability in abundances and levels of sparsity. We use either subsets or taxonomically aggregated versions of *A*_AGP_ to assemble specific scenario count matrices *A*. We then generate synthetic outcomes *y*_*s*_ according to the constrained quadratic log-contrast (qlc) model described in Eq. (8) (see also Fig. 4). In the designed benchmark scenarios we select (uncorrelated) taxa from *A*_AGP_ with different sparsity levels, vary the number of overall features (taxa) *p*, the number and placement of interaction effects 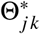,and the noise level *ϵ* ^*^. Given a compositional count matrix *A* (where a pseudocount of 1 is added to all zero counts [8]) and a synthetic outcome *y*_*o*_, we solve the optimization defined in Eq. (12) to estimate the coefficients 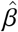 and 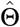 in the sparse quadratic log-contrast models using 5-fold CV. Rather than looking at predictive quality, we measure model quality in terms of estimation error 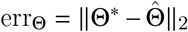 and 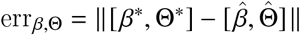, respectively. The error thus indicates how well we can recover the “true” main and interaction effects.

**Fig 4.**
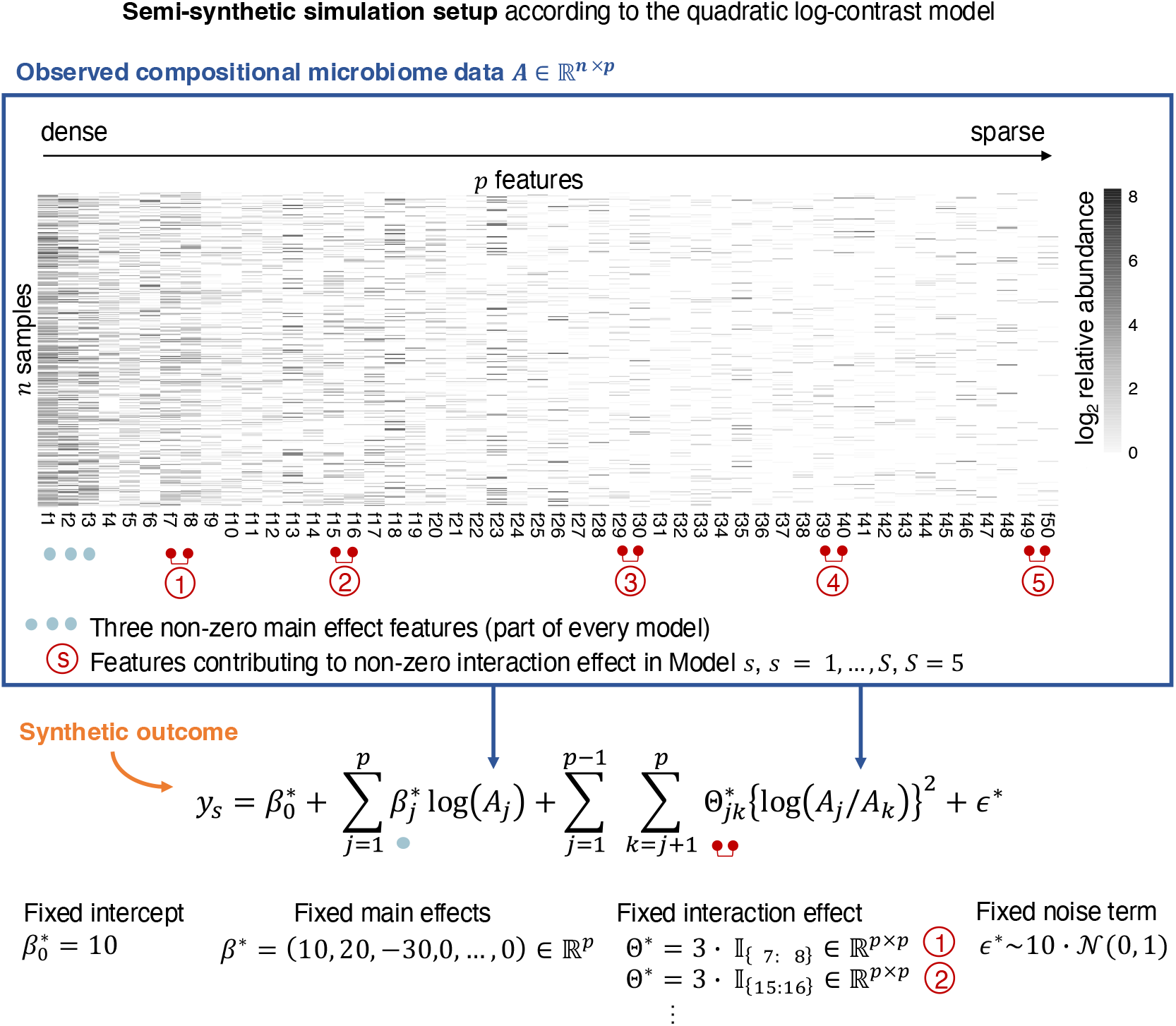
Simulation setup for generating a semi-synthetic outcome *y*_*s*_ for *s* = 1, …, *S* based on the quadratic log-contrast model formulation. Dark blue box: Heatmap of an observed compositional microbiome dataset sorted by sparsity in descending order. Non-zero main effects contributing to each of the *S* = 5 semi-synthetic scenarios (light blue) and features contributing to the non-zero interaction effect in model scenario *s* for *s* = 1, …, *S* (dark red) are highlighted.

#### Feature sparsity influences interaction estimation quality

To test the influence of taxon sparsity on estimation quality, we select a subset of *p* = 50 OTUs from *A*_AGP_ representing a wide range of sparsity levels across a subset of *n* = 300 samples. Figure 4 shows the data matrix *A* and details the generative model for the synthetic outcome *y*_*s*_ for *s* = 1,…, *S* scenarios. In all *S* = 5 scenarios we use a fixed intercept term 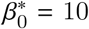,three non-zero main effects (*β*_1_ = 10, *β*_2_ = 20, and *β*_3_ = −30), and a noise term *ϵ* ^*^ ∼ *c*_1_ · 𝒩 (0, 1) with *c*_1_ = 10.

In each scenario, we consider a single interaction effect 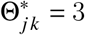 between two features (OTUs) fj and fk. In each scenario, this interaction effect is placed on a different pair of features. In scenario 1, we select feature 7 and 8 (f7:f8) where the product of the features comprises 36% zero entries. In scenario 2, we select feature 15 and 16 (f15:f16) comprising 52% zeros. In scenario 3 we select features 29 and 30 (f29:f30) with 67% zeros, in scenario 4 features 39 and 40 (f39:f40) with 74% zeros, and in scenario 5 features 49 and 50 (f49:f50) with 88% zero entries (see Fig. 4 and Fig. S3b). We also ensure that the selected interaction features are uncorrelated with the main effects (Kendall’s pairwise correlation |τ| *<* 0.2, see also Fig. S3c).

Table 1 summarizes the median and variance of the estimation error err_Θ_ of the sparse quadratic log-contrast model for the five different scenarios over ten simulation replicates. We observe that feature sparsity has a considerable effect on estimation quality of the single 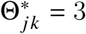 with a marked increase in median estimation error beyond 52% sparsity. Inspection of the solutions across the entire *λ*-path also shows that, while the main effects can be well estimated in all scenarios, the interaction effect cannot be identified anymore for scenarios 4 and 5, respectively, with estimated effect sizes on the same order as the noise level (see Fig. S4). Our simulation results thus give practical guidance when analyzing sparse compositional count data, suggesting that combinations of very sparse features can be *a priori* excluded from the interaction terms.

**Table 1.**
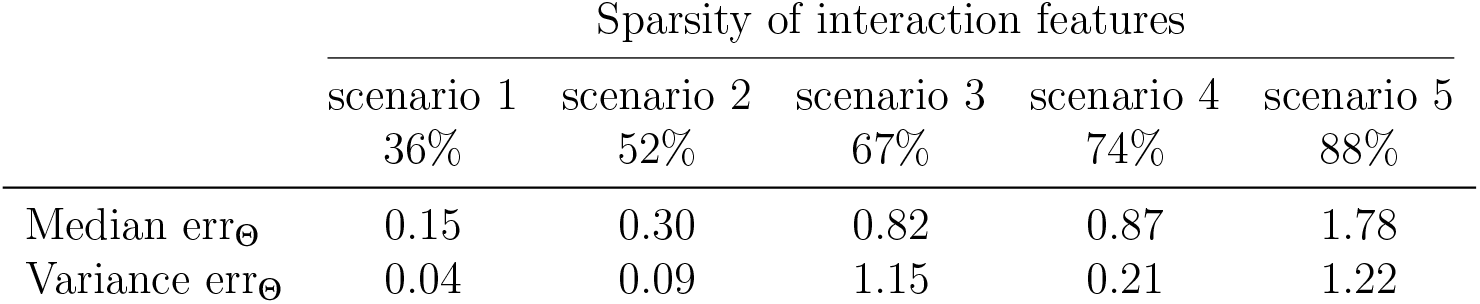
Estimation error 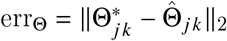 for all five scenarios (10 replicates). Sparsity of interaction features

|  | Sparsity of interaction features |  |  |  |  |
| --- | --- | --- | --- | --- | --- |
|  | scenario 1<br>36% | scenario 2<br>52% | scenario 3<br>67% | scenario 4<br>74% | scenario 5<br>88% |
| Median $\text{err}_{\Theta}$ | 0.15 | 0.30 | 0.82 | 0.87 | 1.78 |
| Variance $\text{err}_{\Theta}$ | 0.04 | 0.09 | 1.15 | 0.21 | 1.22 |

#### Interaction models require strong signals for accurate estimation

To understand the noise dependency of estimating main and interaction effects in quadratic log-contrast models, we next create a low-dimensional benchmark by aggregating OTUs in *A*_AGP_ to the phylum level using the available taxonomic information. We consider the *p* = 10 most prevalent phyla across *n* = 6266 individuals, leading to a dense compositional count matrix 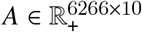.This aggregation ensures that feature sparsity effects on model estimation are not present in the following scenarios. We fix the intercept term at 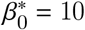,assign six non-zero main effects to the first six features (f1-f6) with *β*^*^ = [10, 20, 30, −10, −20, −30, 0,…, 0]^*T*^ ∈ R^10^, and introduce three non-zero interaction effects (f1:f3, f8:f10, and f9:f10) with 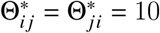 for the respective *ij* pairs and Θ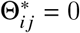 otherwise.

We generate L=8 different scenarios by multiplying the standard normal noise term *ϵ* ^*^ by a constant factor *c*_*l*_, *l* = 1,…, 8, with *c*_*l*_ = {10, 20, 70,100, 200, 300, 400, 500}. Each scenario thus only differs in their noise level 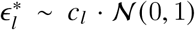For the specified semi-synthetic simulation models, these noise levels correspond to signal-to-noise ratios (SNRs), defined as 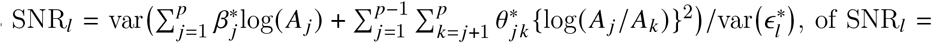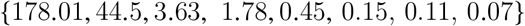.

Table 2 summarizes the median and the variance of the estimation error err_*β*,Θ_ of the sparse qlc models over 10 replicates. We observe remarkable estimation performance when the SNR is high (≥ 44.5). However, the quality deteriorates quickly when the SNR is smaller than one. For the smallest SNR_8_ = 0.07, the median err_*β*,Θ_ of 138.78 is close to an error of 150, achieved by the empty model (i.e., all estimated main and interaction effects are set to 0).

**Table 2.** Median and Variance of the estimation error 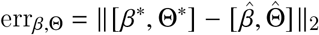 for the eight semi-synthetic scenarios with different signal-to-noise ratios SNR_*l*_.

| $\text{SNR}_l$ | 178.01 | 44.5 | 3.63 | 1.78 | 0.45 | 0.15 | 0.11 | 0.07 |
| --- | --- | --- | --- | --- | --- | --- | --- | --- |
| Median $\text{err}_{\beta, \Theta}$ | 3.89 | 5.07 | 30.21 | 38.44 | 66.96 | 135.46 | 137.34 | 138.78 |
| Variance $\text{err}_{\beta, \Theta}$ | 1.74 | 4.61 | 38.41 | 57.44 | 362.86 | 272.38 | 67.87 | 30.35 |

This suggests that, for low SNRs, neither main nor interaction effects can be disentangled from the noise. This typical behavior of sparse models is also reflected in the error variances which are low both for high SNR scenarios (implying consistent high estimation accuracy) and low SNR scenarios (implying consistent near-empty model estimation).

#### Neglecting interaction effects can lead to model misinterpretation

We next investigated the questions (i) to what extend the inclusion of interaction effects can improve predictive performance compared to the baseline linear log-contrast model and (ii) how model misspecification influences model interpretation. We used a subset of the previous semi-synthetic simulated data SNR_*l*_ = {178.01, 1.78, 0.45, 0.07} and estimated sparse linear log-contrast models (i.e., main effects only) models.

Figure 5 illustrates the behavior and the performance of both linear and quadratic log-contrast models. We first consider the high SNR regime (SNR_1_ = 178.01). Figure 5a shows typical solution paths of both models. As expected, we observe that the interaction model correctly identifies the six main (f1-f6) and three interaction effects (f9:f10, f1:3, f8:f10) at the *λ* values selected by CV (dashed line 1se *λ*_1se_, solid line minimum MSE *λ*_min_). The (misspecified) main effects model, lacking interaction terms, identifies not only the correct main effects but also includes “wrong” main effects. In particular, the model selects f9 early in the solution path with a strong positive effect, followed by a negative effect on f8 and positive effect on f10. This behavior is consistent with the fact that the main effects model compensates for the lack of interactions terms by putting a strong effect on the individual components of the interaction terms f8-f10. Moreover, the misspecification also results in a biased estimation of the f1 and f3 effects (see Fig. 5b), pushing the strong positive effect of f3 close to zero and overestimating the strength of f1. In the absence of a ground truth, the main effects model would thus lead the researcher to a partial misinterpretation of the role of the different features. In the high SNR regime, this misinterpretation is exacerbated by the fact that the main effects model shows excellent predictive performance in terms of *R*^2^ (both train and test *R*^2^ *>* 0.8, see Fig. 5c, panel 1). This effect also persists for lower SNR scenarios (see Fig. 5c, panels 2-4). Nevertheless, the quadratic interaction models, as expected, consistently outperform main effects models in terms of predictive performance by a wide margin of 10-15% even for a medium SNR of 1.78. This performance advantage only vanishes at the lowest SNR of 0.07 (see Fig. 5c, panel 4). Overall, these simulation benchmarks indicate that, in practice, the inclusion of interaction effects compares favorably to the standard linear models both in terms of model identifiability, interpretability, and predictive performance. To test the generality of these results, we also extended the simulation scenarios and analyses for higher-dimensional cases where we aggregated the AGP data to family level (p=50 most prevalent) and included 15-30 main and 20-60 interaction effects. The results of these semi-synthetic scenarios are consistent with the ones described here and are summarized in Fig. S3.

**Fig 5.**
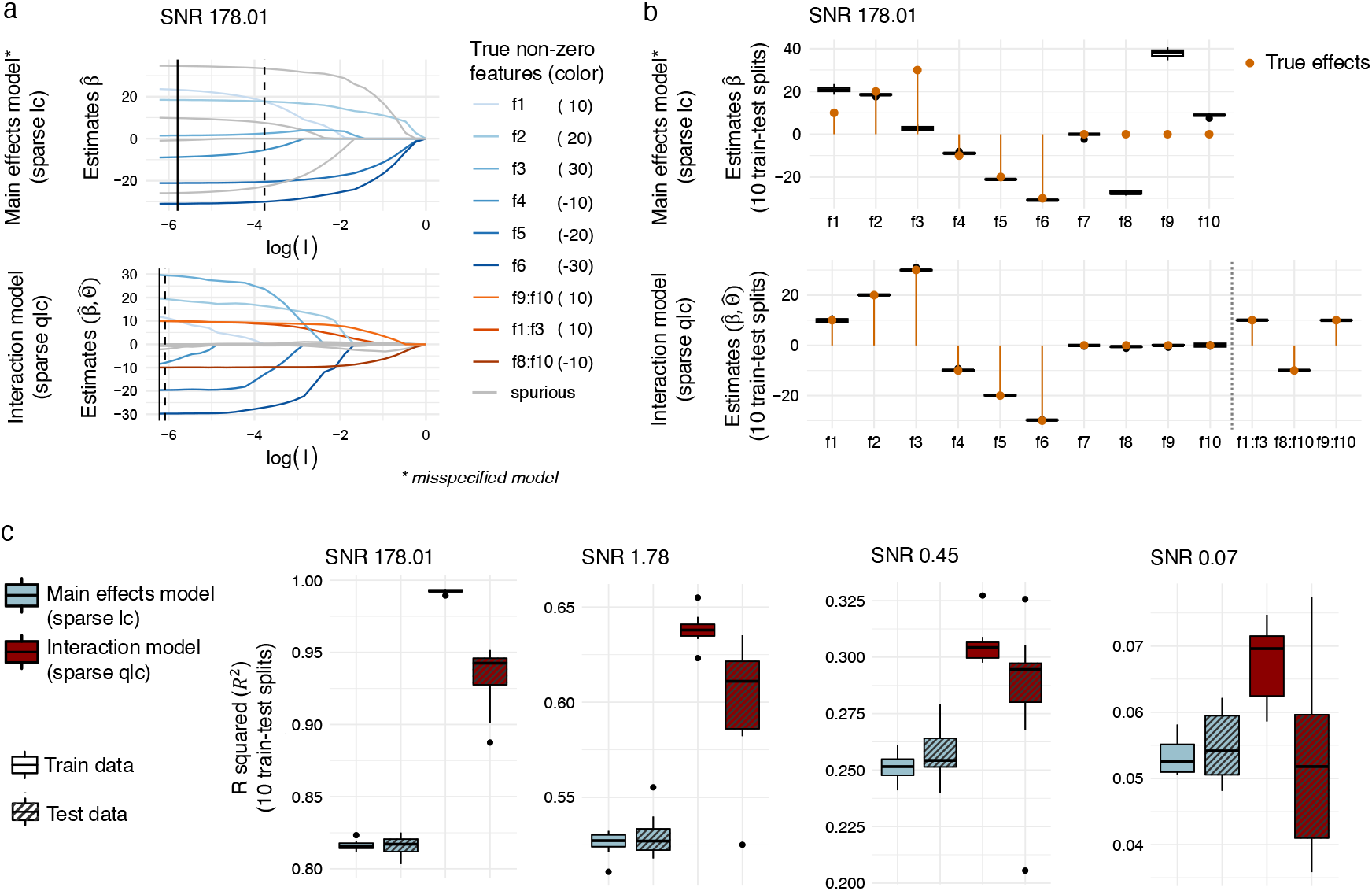
Influence of model misspecification and noise in semi-synthetic scenarios. **a**. Solution path for the misspecified main effects model (sparse lc) and the interaction model (sparse qlc) for the synthetic scenario *l* = 1 with a signal-to-noise ratio (SNR) of 178.01 for one train test split (solid line: *λ*_min_ solution; dashed line: *λ*_1se_ solution). **b**. Estimated coefficients distributions over 10 train test splits corresponding to the solution paths in a. For the interaction model only three non-zero interaction features are shown for visualization purposes. **c**. Comparison of model performance via R squared (*R*^2^) for the main effects model and the interaction effects model on train and test data for four different SNRs.

### Improved global marine salinity prediction from interaction modeling of Tara ocean data

To illustrate the practical benefits of sparse quadratic log-contrast modeling we follow [33] and estimate a marine “community-environment” landscape with ocean salinity concentration as target environmental characteristic from Tara Oceans data [42]. In the original Tara study, temperature was as environmental variable and shown to be well predicted by global microbial community composition. Although this type of modeling is “anti-causal” (i.e., temperature influences microbial abundance rather than the other way around), the good predictability of temperature was taken as evidence for temperature being the “main driver” of microbial diversity in the global ocean. Likewise, it is known that variations in salinity also play a critical role in shaping microbial community diversity [67, 68]. Building the community-salinity landscape thus quantifies how strong of a driver salinity is on shaping microbial composition.

The Tara Oceans data includes relative microbial abundances in form of metagenomic OTUs (mOTUs) [69] of ocean surface water and associated environmental covariates at *n* = 136 sampling sites. We aggregated the *p* = 8916 mOTUs present in the data [33] to family level and selected the *p* = 30 most prevalent families to avoid overly sparse features. We learned both sparse linear and quadratic log-contrast model over ten train-test splits.

Figure 6 summarizes the key results in terms of median train-test *R*^2^, typical out-of-sample predicted salinity concentrations, and median main and interaction effect sizes. We observe that including interaction effects more than doubles predictive accuracy (see Fig. 6a) and predicts salinity concentrations across a larger dynamic range (≈ [33, 38]ppt, see Fig. 6b). Despite considerable variation in the selected interaction models across the ten train-test splits (see Fig. 6c), we identify seven family main effects and eight interaction effects with consistent (non-zero) effect sizes (highlighted in bold in Fig. 6c). The seven families with non-zero median effects sizes include three families from the SAR11 clade (f39, Surface4, Chesapeake-Delaware Bay), two families from the Oceanospirillales order (ZD0405, JLETNPY6), one family from Rickettsiales order (S25593), and the Halomonadacceae family. Consistent with known functionality, Halomonadacceae as halophilic (“salt-loving”) bacteria are positively associated with salinity concentration. Likewise, the family annotated as Chesapeake-Delaware Bay (from the SAR11 clade) has been isolated from estuarine waters with oligohaline and mesohaline conditions [70, 71], implying a negative relationship with salinity. Overall, eleven families are involved in forming the eight consistent log-contrast interactions: unknown families f8, f13, f40, f55, f60, f96, and Chesapeake-Delaware Bay family from the SAR 11 clade, S25593 from Rickettsiales, Oceanospirillaceae, Sphingomonadaceae, Bacteriovoracaceae. The two strongest negative interactions effects are formed by the f55:Sphingomonadaceae and f13:f96 pairs. The two strongest positive interaction effects are formed by the f8:f60 and f40:f96 pairs, respectively. Additionally, we observe the Chesapeake-Delaware Bay:Bacteriovoracaceae pair to be negatively associated with salinity, consistent with the observed main effect of the Chesapeake-Delaware Bay family. Furthermore, the enrichment of SAR 11 clade families in the estimated interaction models suggest a significant habitat diversity within the SAR11 clade with respect to salinity concentrations, reflecting the adaptation mechanisms to specific high-, medium-, and low-saline environments within the lineage [72, 73, 74, 75].

**Fig 6.**
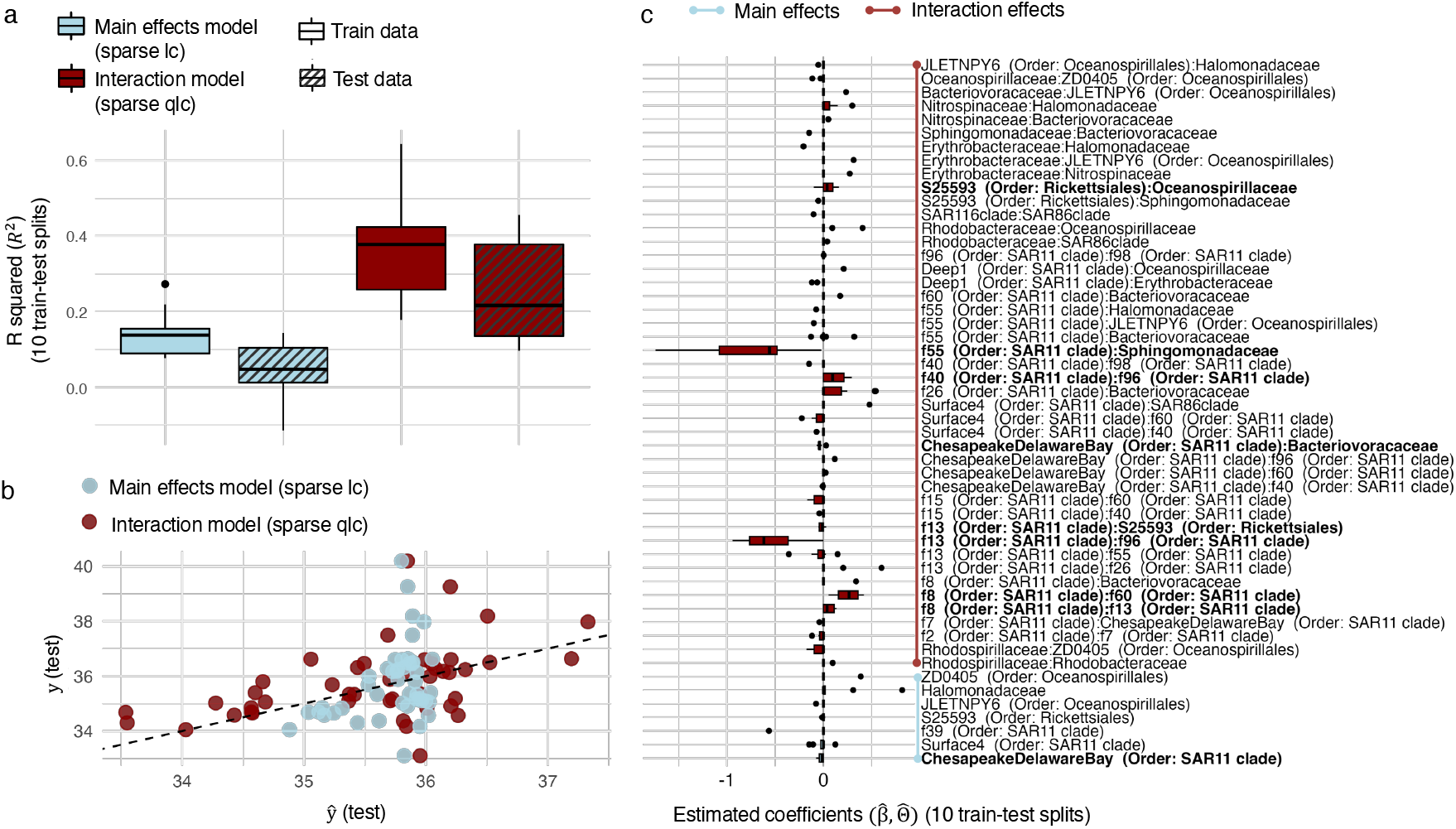
Summary plots of the Tara ocean data on family level for salinity prediction. **a**. Train and test set *R*^2^ distribution over ten train-test splits for the main effects sparse log contrast model (sparse lc) and the sparse quadratic log-contrast model (sparse qlc). **b**. Scatter plot between the observed test set outcome *y* (salinity) and prediction 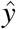 from the sparse lc and the sparse qlc models. **c**. Distribution of estimated main effect and interaction effect coefficients of the sparse qlc models over ten train-test splits. Only features (main or interaction features) with non-zero mean coefficient are shown. Features with a non-zero *median* are bold.

### Microbial data types impact the structure and predictive performance of interaction models

We have thus far studied the behavior and the performance of interaction models individually for the three most prevalent microbial data types: quantitative microbial abundances [21], presence-absence data, and (sequencing-derived) relative abundances. Two natural remaining questions are (i) to what extent the available microbial data type influence interaction model predictive performance *and* (ii) how the structure of the estimated interaction models changes with microbial data type, thus influencing model interpretation.

To shed light on these questions, we revisit the ARG prediction task using the MetaCardis cohort data [41]. We consider the *p* = 30 most abundant genera and transform the original absolute abundance data into presence-absence and relative abundance data, respectively.

This enables us to study the same prediction task of estimating the number of ARGs from genus compositions while simultaneously analyzing the structure of the resulting genus main and genus-genus interaction effects across the different data types.

We next estimated the respective data-type specific sparse quadratic interactions models (without hierarchy constraints) from the three data sets, using identical train-test splits and 5-fold CV for model selection. We repeated the experiment on ten different train-test splits and report predictive performance using median test-set *R*^2^ and median effect sizes.

Figure 7 summarizes the key results of the experiments. We observe a consistent drop in predictive performance with decreasing information content of the data. We achieve the best median *R*^2^ = 0.53 for absolute abundance data, followed by an *R*^2^ = 0.44 for relative abundance data, and an *R*^2^ = 0.28 for genus presence-absence data. Figure 7a illustrates observed vs. predicted ARG numbers on a representative test set for each data type, observing a consistent decrease in overall correlation.

**Fig 7.**
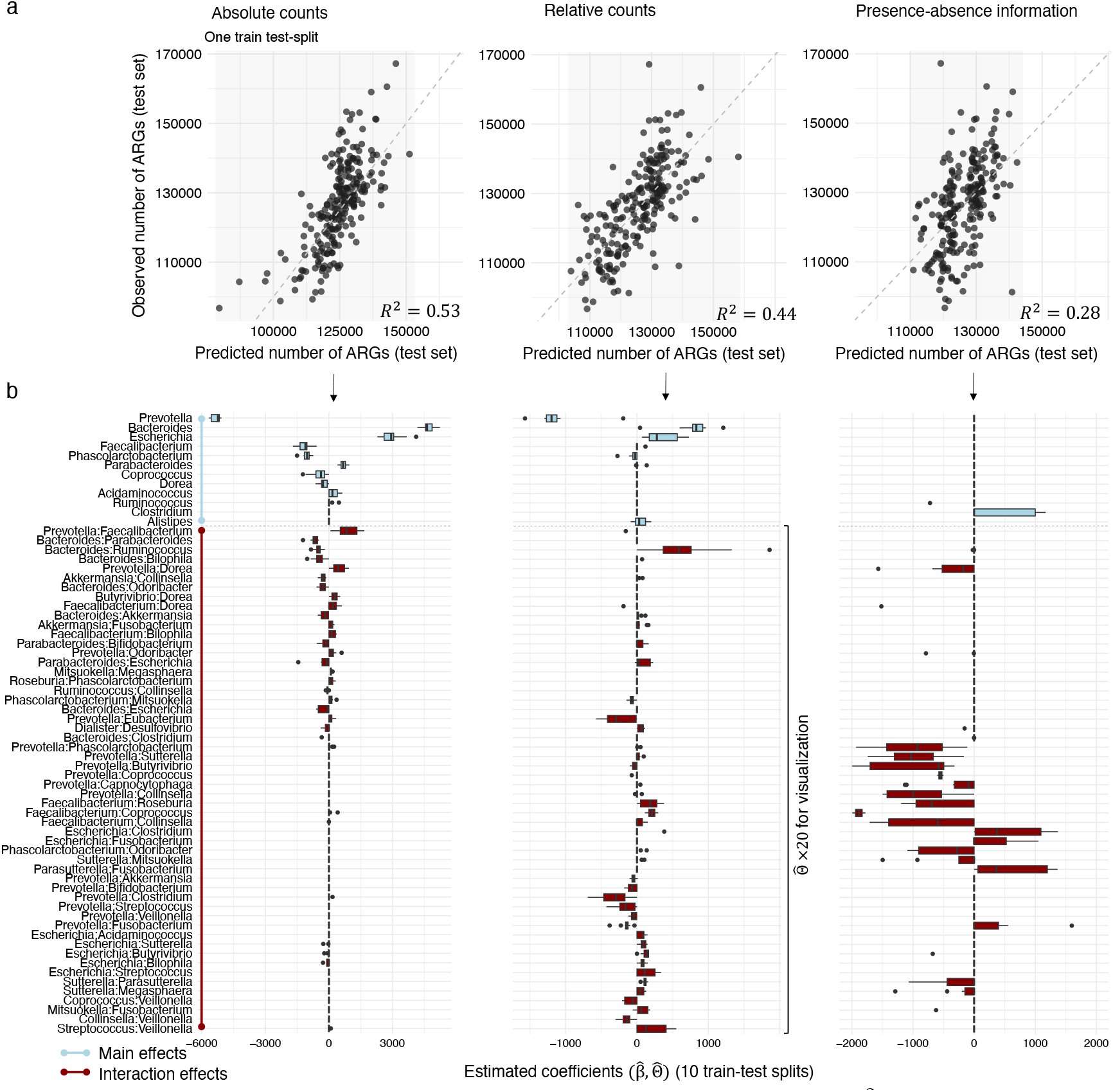
a. Scatter plots (one train test split) and test set R squared *R*^2^ (average over 10 train test splits) for the comparison of the predicted number of anti microbial resistance genes (ARGs) and the observed number of ARGs on a test dataset based on the absolute count information of genera, the relative count information of genera and the presence-absence information of genera of the MetaCardis cohort. **b**. Distribution of coefficients over 10 train test splits for the superset of coefficients that are non-zero in at least one of the three data modalities.

To analyze the corresponding interaction model structures, we focus on the union of all main and interaction effects whose medians were non-zero in at least one of the three interaction models. Figure 7b summarizes the distribution of the effect sizes for both main (light blue) and interaction effects (red) of the estimated interaction models. Overall, we observe pronounced differences in the structure of the different interaction models. While we see agreement in the three strongest main effects of the absolute and relative abundance models (positive effects of Bacteroides and Escherichia and negative effects of Prevotella, respectively), other absolute abundance-based main effects are barely present in the relative abundance-based and the presence-absence models. Strikingly, the presence -absence models is the only model that posits a high positive main effect on the Chlostridium genus. We observe considerable variation in the remaining interactions. While the absolute abundance-based interaction model estimates positive effects on the Prevotella:Faecalibacterium and Prevotella:Dorea interaction pairs, the relative abundance-based model estimates a large positive effect for the Bacteroides:Ruminococcus ratio and a large negative one on the Prevotella:Clostridium ratio. The presence-absence interaction model comprises many negative interaction effects involving the Prevotella genus while lacking a Prevotella main effect. This is consistent with our previous observation that interaction models without hierarchy constraints can make model interpretation challenging.

## Discussion

Learning predictive and interpretable models that relate microbial abundance data to community function, host-associated phenotypes, or environmental covariates is a cornerstone in microbiome data analysis and microbial ecology. To this end, we have introduced a general statistical framework for sparse quadratic interaction modeling that can accommodate all common microbial data types, including sequencing-derived relative abundance data. Our framework unifies several disjoint approaches in the microbial ecology and microbiome data science literature, tying together the recent ecological concept of “community-function landscape” [15, 16, 76, 25] with high-dimensional statistics and compositional data analysis approaches [27, 28, 33].

Using a wide range of application scenarios, including synthetic microbial communities, human gut, and marine microbiome surveys, we have illustrated the practical benefits of applying modern statistical concepts such as hierarchical constraints [36, 37, 38] and stability-based model selection [39, 40] for interpretable predictive modeling. For instance, we could demonstrate that the number of antimicrobial resistance genes (ARGs) in the MetaCardis cohort can be well predicted by genus-level interaction models, extending and improving on prior work on enterotype-based adjustments [60]. Since our estimated genus-based models are both applicable to metagenomics and amplicon sequencing data, future work may include *de novo* application of the ARG prediction models to the large number of available amplicon gut microbiome datasets. This could potentially reveal significant differences in predicted ARG abundances across cohorts and, in turn, point toward hidden confounding.

On the Clark et al. [18] synthetic community dataset, our framweork was able to identify the negative association between *A. caccae* and *D. piger* as the *only* stable empirical interaction that is relevant for the community function of butyrate production. This clarifies not only previous less clear-cut results [18, 76, 15] but also suggests that our framework may prove useful in identifying and designing new synthetic communities with specific community function [77, 15, 78].

For the arguably most important use case of modeling amplicon sequencing data, we have introduced a series of semi-synthetic simulation scenarios that quantified the relationship between prediction and model estimation quality, extreme sparsity of the underlying data, noise corruption, and model misspecification. Together with our study of the community-salinity relationship from the Tara ocean data, these scenarios have provided a first glimpse into the benefits and limitations of using interaction models on high-dimensional relative microbial abundance data. Future applications of this framework will be required to more fully assess its effectiveness and generality both on standardized benchmarks [79, 33] and relevant disease prediction tasks [80].

Finally, our comparative study of using absolute abundance, relative abundance, and presence-absence data for ARG prediction revealed noticeable differences in prediction quality and interaction model structure, highlighting the importance of the available microbial community data type. This interplay between microbial data type and statistical model structure and quality has been reported in related studies. For instance, Vandeputte et al. [21] observed that empirical species-species correlations strongly differ when estimated from sequencing-based quantitative vs. relative abundance data. While a large part of these differences can be ameliorated by using proper statistical correlation methods [10, 81, 82], recent work has also shown that predictive models for disease-microbe associations are strongly confounded when neglecting quantitative information in form of fecal microbial load [83]. We posit that, with the increasing availability of quantitative microbiome data, our framework enables principled comparative main effect and interaction modeling across data types to identify context-*and* data type-dependent (confounded) microbial predictors.

Despite the wide applicability and generality of the presented framework there exist a number of possible improvements and extensions. Firstly, with the expected availability of larger data sets, it is conceivable to extent our statistical framework to higher-order interactions [76] or non-parametric, non-linear interactions [84]. How to effectively deliver higher-order hierarchical and stable interactions in this context is, however, an open problem. Secondly, throughout this manuscript we reported median test performance of the interaction models across multiple train-test split, enabling a more realistic view on prediction performance and its variability. This idea can be more rigorously assessed using the concept of (conformal) prediction sets [85]. Including recent work on prediction sets for sparse interaction models in our statistical framework would potentially further increase the confidence of the practitioner in the quality of the estimated models. Lastly, the focus of this manuscript has been on interaction modeling using microbial abundance data alone. However, when more than one data modality are available, e.g., microbial abundances and additional host or environmental covariates, multimodal interaction modeling extensions are likely to improve predictive performance. While conceptually straightforward, future research needs to assess how to properly normalize and transform different multimodal data types separately or jointly for effective interaction modeling.

In summary, we have introduced statistically principled and reproducible workflows for predictive microbial interaction modeling, available as reproducible R code at https://github.com/marastadler/Microbial-Interactions. We anticipate that these workflows will serve as a baseline for more sophisticated modeling endeavors, delivering new hypotheses and predictions of relevance in microbial community ecology and microbiome research.

## Supporting Information Legends

### Mathematical Models and Equations

- **Quadratic log-ratio model**: Description of the quadratic log-ratio model for compositional data, including equations (S.1) and (S.2), and its extension to the quadratic log-ratio interaction model (qlr).
- **Sparse alr transformed quadratic model**: Explanation of the loss function and optimization problem for the sparse alr transformed quadratic model, as defined in equation (S.3).
- **Sparse quadratic log-ratio model**: Description of the loss function and optimization problem for the sparse quadratic log-ratio model (qlr), as defined in equation (S.4).
- **Comparison of interaction models**: Discussion of three mathematically equivalent models for modeling quadratic interactions with relative microbiome data: (a) the alr transformed quadratic model, (b) the quadratic log-contrast model, and (c) the quadratic log-ratio model.
- **Binary encoding of covariates**: Explanation of the impact of binary encoding (e.g., {0, 1} vs. {−1, 1}) on the interpretation of model coefficients in regression models, including transformations between encodings.

### Figures

- **Figure S1.** Low-dimensional representation (UMAP) of the microbial abundance data *A*^*n*×*p*^ with *n* = 690 individuals (corresponding to the number of points) and *p* = 30 most prevalent genera. **a**. UMAP representation colored by the number of ARGs. **b**. UMAP representation with enterotypes highlighted, indicating a positive correlation of ARGs with *Bacteroides 1*, a particularly strong positive association of ARGs with *Bacteroides 2*, and a negative association of ARGs with *Prevotella*.
- **Figure S2.** Distribution of estimated coefficients with non-zero median over 10 train test splits in the sparse linear model.
- **Figure S3.** Semi-synthetic simulation setup for varying feature sparsity levels. **a**. Simulation setup for generating a synthetic outcome *y*_*s*_ for *s* = 1, …, *S* based on the quadratic log-contrast model formulation. **b**. Heat map of the OTU table carrying compositional information for a subset of *p* = 50 OTUs from the American Gut cohort sorted by sparsity in descending order. Non-zero main effects contributing to each of the *S* = 5 semi-synthetic scenarios (light blue) and features contributing to the non-zero interaction effect in model scenario *s* for *s* = 1, …, *S* (dark red) are highlighted. **c**. Kendall’s pairwise correlations τ between features that have non-zero effects in the models *s* = 1, …, *S*. They should be as uncorrelated as possible (|τ| *<* .2) to eliminate effects of correlated features.
- **Figure S4.** Solution path of the interaction model (sparse qlc) for the *S* = 5 semi-synthetic simulation setups for varying feature sparsity levels.
- **Figure S5.** Semi-synthetic data simulations according to the sparse quadratic log-contrast model based on the American Gut Project data at the family level from *A*_AGP_, comprising *n* = 6266 samples and the *p* = 50 most prevalent families, leading to a compositional count matrix 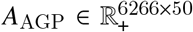We fix the intercept term at 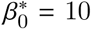 and vary the number of non-zero main and interaction effects. The non-zero entries are sampled from a normal distribution *β*^*^ ∼ *𝒩* (0, σ) and Θ^*^ ∼ 𝒩 (0, σ). Panels (a, b, and c) show the distributions of the estimated coefficients across 10 train-test splits, highlighting the effect of varying the number of 15 to 30 non-zero main effects (a: 15, b: 20, c: 30) and 20 to 60 interaction coefficients (a: 20, b: 30, c: 60).

