## Supplementary Material for "Predictive modeling of microbial data with interaction effects"

### Supplementary information

#### Quadratic log-ratio model

Another way of accounting for compositionality in regression models is to build log-ratios between all possible pairs of features in $A \in \mathbb{R}_+^{n \times p}$. This approach is referred to the (all-pairs) log-ratio model [31], which is given by

$$y = \beta_0 + \sum_{j=1}^{p-1} \sum_{k=j+1}^{p} \beta_{j,k} \log(A_j/A_k) + \epsilon, \tag{S.1}$$

where the main effect coefficient $\beta_{j,k}$ corresponds to the pairwise (log-ratio) effect of $A_j$ and $A_k$.

In the same way as in 8 the log-ratio model can be extended to a quadratic version, the quadratic log-ratio interaction model (qlr), namely,

$$y = \beta_0 + \sum_{j=1}^{p-1} \sum_{k=j+1}^{p} \beta_{j,k} \log(A_j/A_k) + \frac{1}{2} \sum_{j \neq k} \Theta_{jk} \log(A_j/A_k)^2 + \epsilon, \tag{S.2}$$

where the main effect coefficient $\beta_{j,k}$ corresponds to the pairwise (log-ratio) effect of $A_j$ and $A_k$, with $\beta \in \mathbb{R}^{p(p-1)/2}$ and the interaction effect coefficient $\Theta_{jk}$ corresponds to the quadratic (log-ratio) effect of $A_j$ and $A_k$, with $\Theta = \Theta^T \in \mathbb{R}^{p \times p}$. There exists a linear transformation between the main effect coefficients $\beta_j$ in model 7 and model 8 and the main effects coefficients $\beta_{j,k}$ in model S.1 and model S.2, $\beta_j = -\sum_{k=1}^{j-1} \beta_{k,j} + \sum_{k=j+1}^{p} \beta_{j,k}$, implying that the zero-sum constraint on $\beta \in \mathbb{R}^p$ is inherently met in the linear and quadratic log-ratio model. While the models are mathematically equivalent, their interpretations are different and the choice might depend on the particular data application.

#### Sparse alr transformed quadratic model

Given $A \in \mathbb{R}^{n \times p}$ is a matrix containing the relative abundance information of $p$ microbial taxa, the loss function in 9 for the sparse alr transformed quadratic model, introduced in 6, is defined as

$$l^{\text{qalr}}(\beta_0, \beta, \Theta) = \left\| y - \beta_0 - \sum_{j=1}^{p-1} \beta_j C_j + \frac{1}{2} \sum_{j=1}^{p-1} \sum_{k=1}^{p-1} \Theta_{jk} C_j C_k \right\|_2^2,$$

with $C_j = \log(A_j/A_p)$, $j = 1, \ldots, p-1$. The model does not require further constraints on the model parameters, such that $c(\beta_0, \beta, \Theta) = \emptyset$. Consequently, the optimization problem is given by

$$\underset{\beta_0, \beta, \Theta}{\text{minimize}}\, \rho(l^{\text{qalr}}, \beta_0, \beta, \Theta). \tag{S.3}$$

In the linear model case the loss function in the optimization problem reduces to $l^{\text{alr}}(\beta_0, \beta) = \left\| y - \beta_0 - \sum_{j=1}^{p-1} \beta_j C_j \right\|_2^2$.

#### Sparse quadratic log-ratio model

The loss function of the sparse quadratic log-ratio (qlr) model corresponding to the interaction model for compositional data, introduced in S.2, is defined as

$$l^{\text{qlr}}(\beta_0, \beta, \Theta) = \frac{1}{2} \left\| Y - \beta_0 - \sum_{j=1}^{p-1} \sum_{k=j+1}^{p} \beta_{j,k} \log(A_j/A_k) - \frac{1}{2} \sum_{j \neq k} \Theta_{jk} \log(A_j/A_k)^2 \right\|_2^2.$$

This model does not require further constraints on the model parameters, such that $c(\beta_0, \beta, \Theta) = \emptyset$. The optimization problem for the sparse quadratic log-ratio model is therefore given as

$$\underset{\beta_0, \beta, \Theta}{\text{minimize}}\, \rho(l^{\text{qlr}}, \beta_0, \beta, \Theta). \tag{S.4}$$

In the sparse log-ratio model, that is linear in the features and corresponds to the model in S.1, the loss function reduces to $l^{\text{lr}}(\beta_0, \beta) = \frac{1}{2} \left\| Y - \beta_0 - \sum_{j=1}^{p-1} \sum_{k=j+1}^{p} \beta_{j,k} \log(A_j/A_k) \right\|_2^2$. The $p(p-1)/2$-dimensional sparse log-ratio problem has been shown to be equivalent to the sparse log-contrast model problem for $\lambda^{\text{qlr}} = 2\lambda^{\text{qlc}}$ [31]. This equality can be directly translated to their quadratic extensions. As the dimensionality of the predictor space in the $p(p-1)/2$-dimensional log-ratio model becomes computationally inefficient for large $p$, the authors in [31] propose a two-stage procedure that involves a pre-selection step for covariates to reduce the predictor space before applying the log-ratio lasso. This two-step procedure can be directly applied to the $2 \cdot p(p-1)/2$-dimensional quadratic log-ratio lasso, introduced in S.2, in scenarios where $p$ is large.

#### Comparison and interpretation of different interaction models for relative microbiome data

We introduce three mathematically equivalent ways of modeling quadratic interactions with relative input data: (a) the alr transformed quadratic model, (b) the quadratic log-contrast model, and (c) the quadratic log-ratio model. The three models differ in terms of interpretability, dimensionality, and optimization. In the $\tilde{p} + \tilde{p}(\tilde{p}-1)/2$-dimensional alr transformed quadratic model, where $\tilde{p} = p - 1$, the interpretation of coefficients is consistently tied to a reference feature, which is enforced to be part of the model in the regularized model case. It is important to note, as highlighted by [45], that the choice of this reference feature can greatly influence the interpretability and applicability of the model results. Selecting a reference that is consistent across samples or enhances isometric properties can optimize the interpretability, making the model outcomes more robust and meaningful. This model can be optimized without constraints on the main effects and can be flexibly extended to hierarchical interactions. The $p+p(p-1)/2$-dimensional quadratic log-contrast model requires a zero-sum constraint on the main effects but offers a more convenient expression that does not require the assignment of a reference feature. In this model, each main effect is interpreted with respect to all other features. The $2 \cdot p(p-1)/2$-dimensional quadratic log-ratio model does not require a constraint on the main effect coefficients in the optimization, as this property is automatically met, and allows for the interpretation of the relative effects between all pairs of features. Notably, the interpretation of the interaction coefficients in both the quadratic log-contrast model and the quadratic log-ratio model are identical. Although the log-contrast model in the linear case enjoys the clear advantage of lower dimensionality compared to the log-ratio model, this distinction becomes less relevant when introducing interactions. Hence, the primary criterion for selecting between the models should be based on the preferred interpretation.

#### Binary encoding of covariates in regression models

Whether to encode the input data in a regression model as $B \in \{0,1\}^{n \times p}$ or as $B \in \{-1,1\}^{n \times p}$ has an impact on the interpretations of the model coefficients. In the main effects model case with $p = 1$, for $B \in \{0,1\}^{n \times p}$, the outcome $y$ would be given by

$$y = \begin{cases} \beta_0 & \text{if } B_1 = 0 \\ \beta_0 + \beta_1 & \text{if } B_1 = 1. \end{cases}$$

The interpretation of $\beta_0$ in this case is the effect of 'absence' and the interpretation of $\beta_1$ is the difference between the effect of 'presence' and the effect of 'absence'. The interaction model, illustrated for $p = 2$, is given by

$$y = \begin{cases} \beta_0 & \text{if } B_1 = 0 \text{ and } B_2 = 0 \\ \beta_0 + \beta_1 & \text{if } B_1 = 1 \text{ and } B_2 = 0 \\ \beta_0 + \beta_2 & \text{if } B_1 = 0 \text{ and } B_2 = 1 \\ \beta_0 + \beta_1 + \beta_2 + \Theta_{12} & \text{if } B_1 = 1 \text{ and } B_2 = 1. \end{cases}$$

Here, $\beta_0$ is the effect of co-absence, and $\beta_j$ is the effect of the difference of co-absence and presence of $B_j$, for $j = 1,2$. The interaction term $\Theta_{12}$ is the additional effect when both features are 1.
If $B \in \{-1,1\}^{n \times p}$, the outcome $y$ in the main effects model is given by

$$y = \begin{cases} \beta_0 - \beta_1 & \text{if } B_1 = -1 \\ \beta_0 + \beta_1 & \text{if } B_1 = 1. \end{cases}$$

The interpretation here is that $\beta_0$ is the mean effect of the two group means, and $2\beta_1$ is the difference of the two conditions in mean.
In the interaction model $y$ is given by

$$y = \begin{cases} \beta_0 - \beta_1 - \beta_2 + \Theta_{12} & \text{if } B_1 = -1 \text{ and } B_2 = -1 \\ \beta_0 + \beta_1 - \beta_2 - \Theta_{12} & \text{if } B_1 = 1 \text{ and } B_2 = -1 \\ \beta_0 - \beta_1 + \beta_2 - \Theta_{12} & \text{if } B_1 = -1 \text{ and } B_2 = 1 \\ \beta_0 + \beta_1 + \beta_2 + \Theta_{12} & \text{if } B_j = 1 \text{ and } B_k = 1. \end{cases}$$

Now, $\beta_0$ represents the mean of the four group means (if the design is completely balanced this is the overall mean). The main effect coefficients $\beta_j$, $j \in \{1, 2\}$ are the average difference effects between the two conditions the respective feature can take. The interaction effect $2\Theta_{jk}$ explains the difference between the two conditions when either both features are present or absent and when only one of the two features is present.
Moreover, there exists a linear transformation between the coefficients of both encodings. We denote all coefficients in the 0 and 1 encoding as $\tilde{\beta}$ and the coefficients in the -1 and 1 encoding as $\beta$. The transformation between both encodings in the quadratic interaction model for $p = 2$ is given by

$$\begin{aligned} \tilde{\beta}_0 &= \beta_0 - \beta_1 - \beta_2 + \Theta_{12} \\ \tilde{\beta}_1 &= 2(\beta_1 - \Theta_{12}) \\ \tilde{\beta}_2 &= 2(\beta_2 - \Theta_{12}) \\ \tilde{\Theta}_{12} &= 4\Theta_{12}. \end{aligned}$$

For $p \geq 2$ this can be translated to a general form as by

$$\begin{aligned} \tilde{\beta}_0 &= \beta_0 - \sum_{j=1}^{p} \beta_j + \sum_{j=1}^{p-1} \sum_{k=j+1}^{p} \Theta_{jk} \\ \tilde{\beta}_j &= 2\beta_j - 2 \sum_{\substack{k=1 \\ k \neq j}}^{p} \Theta_{jk}, \text{ for } j = 1, ..., p \\ \tilde{\Theta}_{jk} &= 4\Theta_{jk}, \text{ for } j = 1, ..., p-1, \; k = j+1, ..., p. \end{aligned}$$

The transformation from one encoding to the other can be derived by replacing the input matrix in the model according to this equation: $B^{\{-1,1\}} = 2B^{\{0,1\}} - 1$.

#### Alr transformed model versus constrained log contrast model

Main effects only:

$$ALR = \sum_{j=1}^{p-1} \beta_j \log \frac{X_j}{X_p} = \sum_{j=1}^{p-1} \beta_j \log X_j - \left( \sum_{j=1}^{p-1} \beta_j \right) \log X_p$$

Thus, we can define $\beta_p := -\sum_{j=1}^{p-1} \beta_j$ and then write this as $\sum_{j=1}^{p} \beta_j \log X_j$ s.t. $\sum_{j=1}^{p} \beta_j = 0$,

which corresponds to the log contrast model.

Model with interactions:

$$\begin{aligned}
ALR &= \sum_{j=1}^{p-1} \beta_j \log \frac{X_j}{X_p} + \sum_{1 \le j,k \le p} \Theta_{jk} \log \frac{X_j}{X_p} \log \frac{X_k}{X_p} \\
&= \sum_{j=1}^{p-1} \beta_j \log X_j + \left( -\sum_{j=1}^{p-1} \beta_j \right) \log X_p \\
&+ \sum_{1 \le j < k < p} \Theta_{jk} \left( \log X_j \log X_k - \log X_j \log X_p - \log X_k \log X_p + \log^2 X_p \right) \\
&= \sum_{j=1}^{p} \beta_j \log X_j \quad (\text{taking } \sum_{j=1}^{p} \beta_j := 0) \\
&+ \sum_{1 \le j,k \le p} \Theta_{jk} \log X_j \log X_k - 2 \log X_p \sum_{j=1}^{p-1} \left( \sum_{k=1}^{p-1} \Theta_{jk} \right) \log X_j + \left( \sum_{j,k} \Theta_{jk} \right) \log^2 X_p
\end{aligned}$$

For $j < p, \Theta_{jp} = \Theta_{pj} := -\sum_{k=1}^{p-1} \Theta_{jk}$

For $\Theta_{pp} := \sum_{1 \le j,k \le p} \theta_{jk}$.

Note: $\sum_{j=1}^{p} \Theta_{jp} = \sum_{j=1}^{p-1} \left( -\sum_{k=1}^{p-1} \Theta_{jk} \right) + \left( \sum_{1 \le j,k < p} \Theta_{jk} \right) = 0$

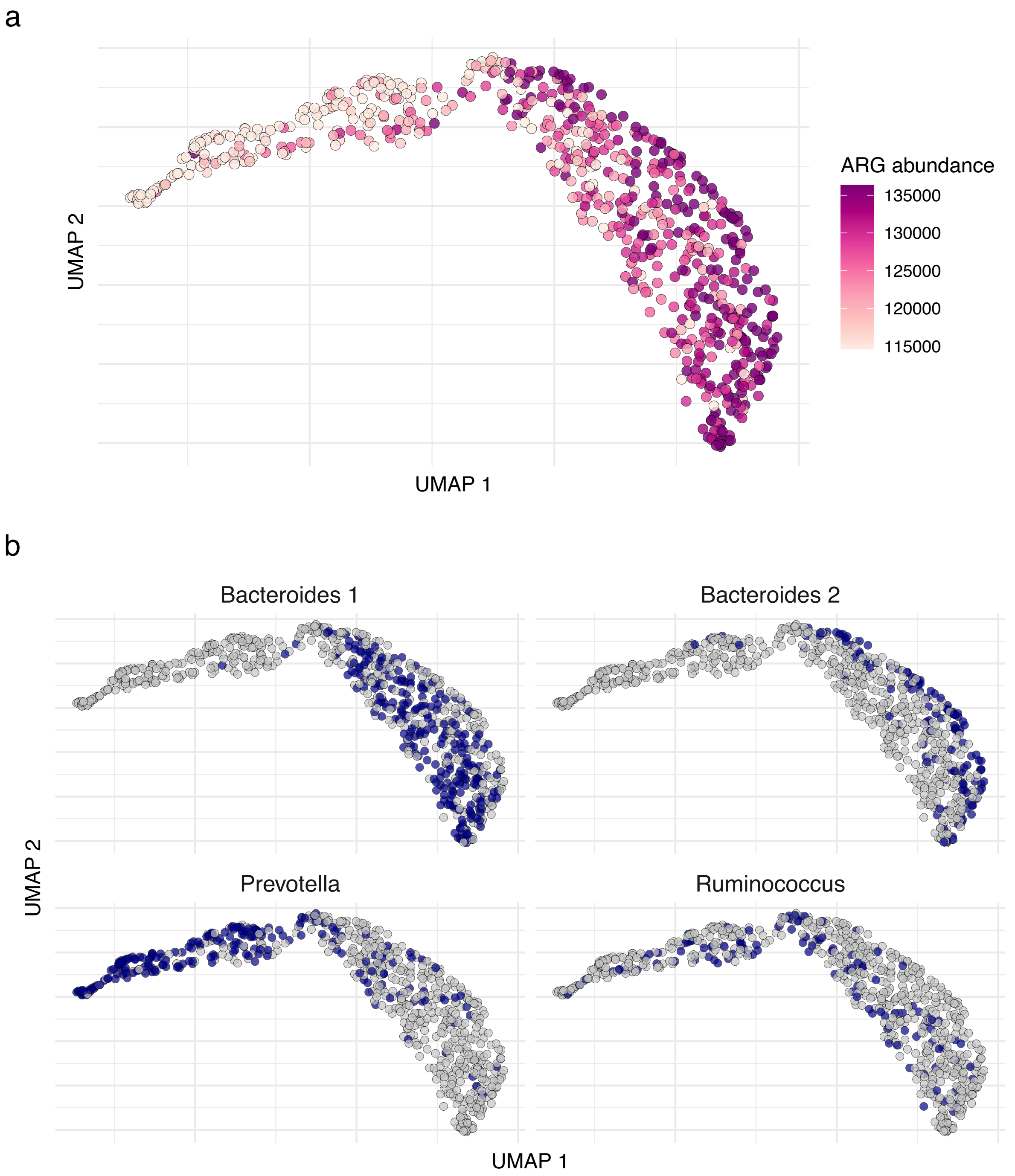


**Fig S1.** Low-dimensional representation (UMAP) of the microbial abundance data $A^{n \times p}$ with $n = 690$ individuals (corresponding to the number of points) and $p = 30$ most prevalent genera. **a.** UMAP representation colored by the number of ARGs. **b.** UMAP representation with enterotypes highlighted, indicating a positive correlation of ARGs with *Bacteroides 1*, a particularly strong positive association of ARGs with *Bacteroides 2*, and a negative association of ARGs with *Prevotella*.

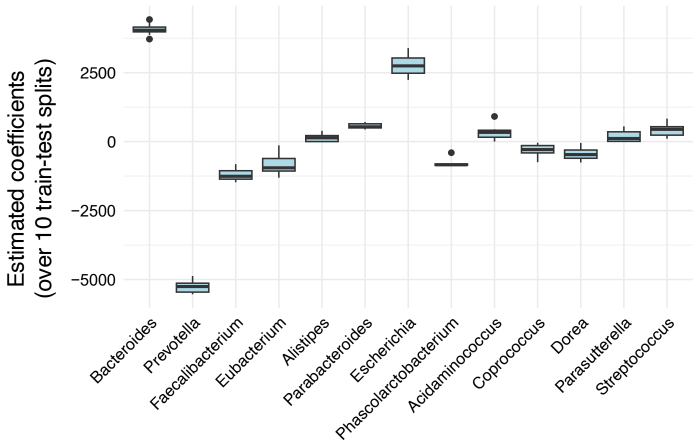


**Fig S2.** Distribution of estimated coefficients with non-zero median over 10 train test splits in the sparse linear model.

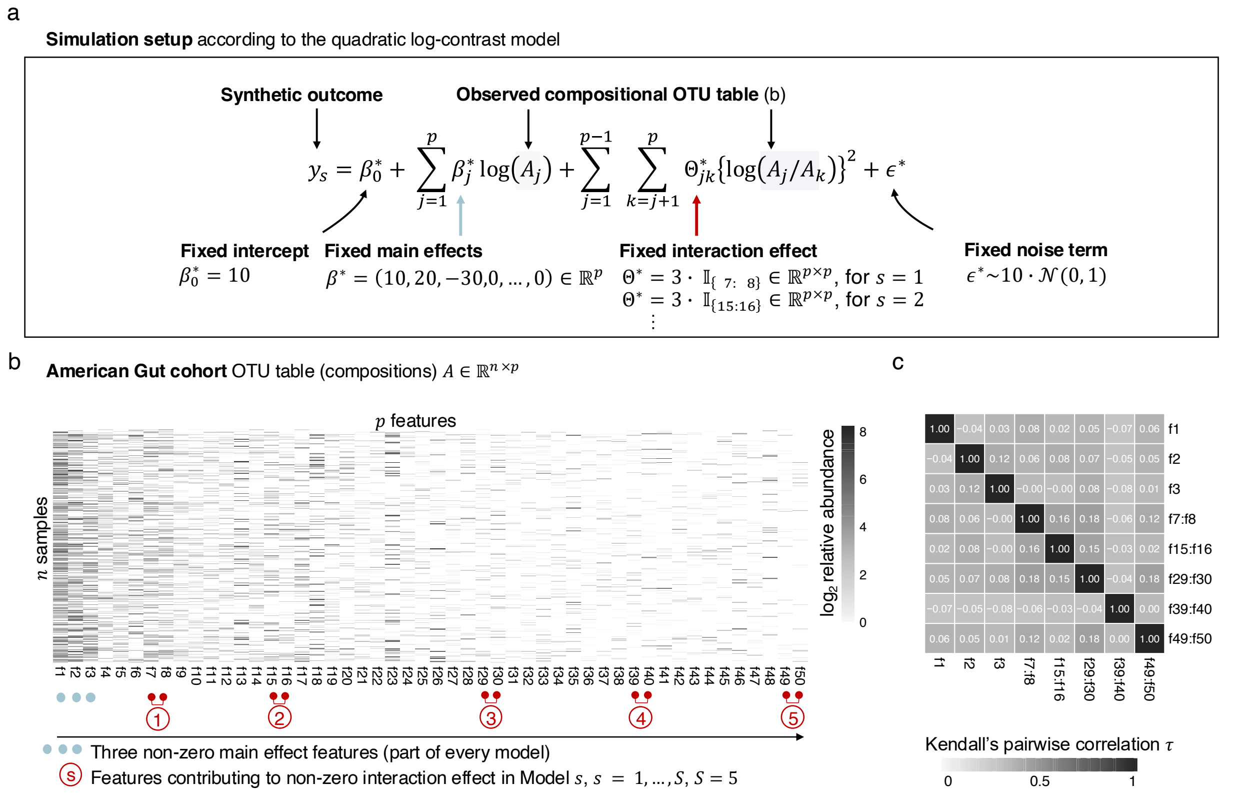


**Fig S3.** Semi-synthetic simulation setup for varying feature sparsity levels. **a.** Simulation setup for generating a synthetic outcome $y_s$ for $s = 1, \ldots, S$ based on the quadratic log-contrast model formulation. **b.** Heat map of the OTU table carrying compositional information for a subset of $p = 50$ OTUs from the American Gut cohort sorted by sparsity in descending order. Non-zero main effects contributing to each of the $S = 5$ semi-synthetic scenarios (light blue) and features contributing to the non-zero interaction effect in model scenario $s$ for $s = 1, \ldots, S$ (dark red) are highlighted. **c.** Kendall's pairwise correlations $\tau$ between features that have non-zero effects in the models $s = 1, \ldots, S$. They should be as uncorrelated as possible ($|\tau| < .2$) to eliminate effects of correlated features.

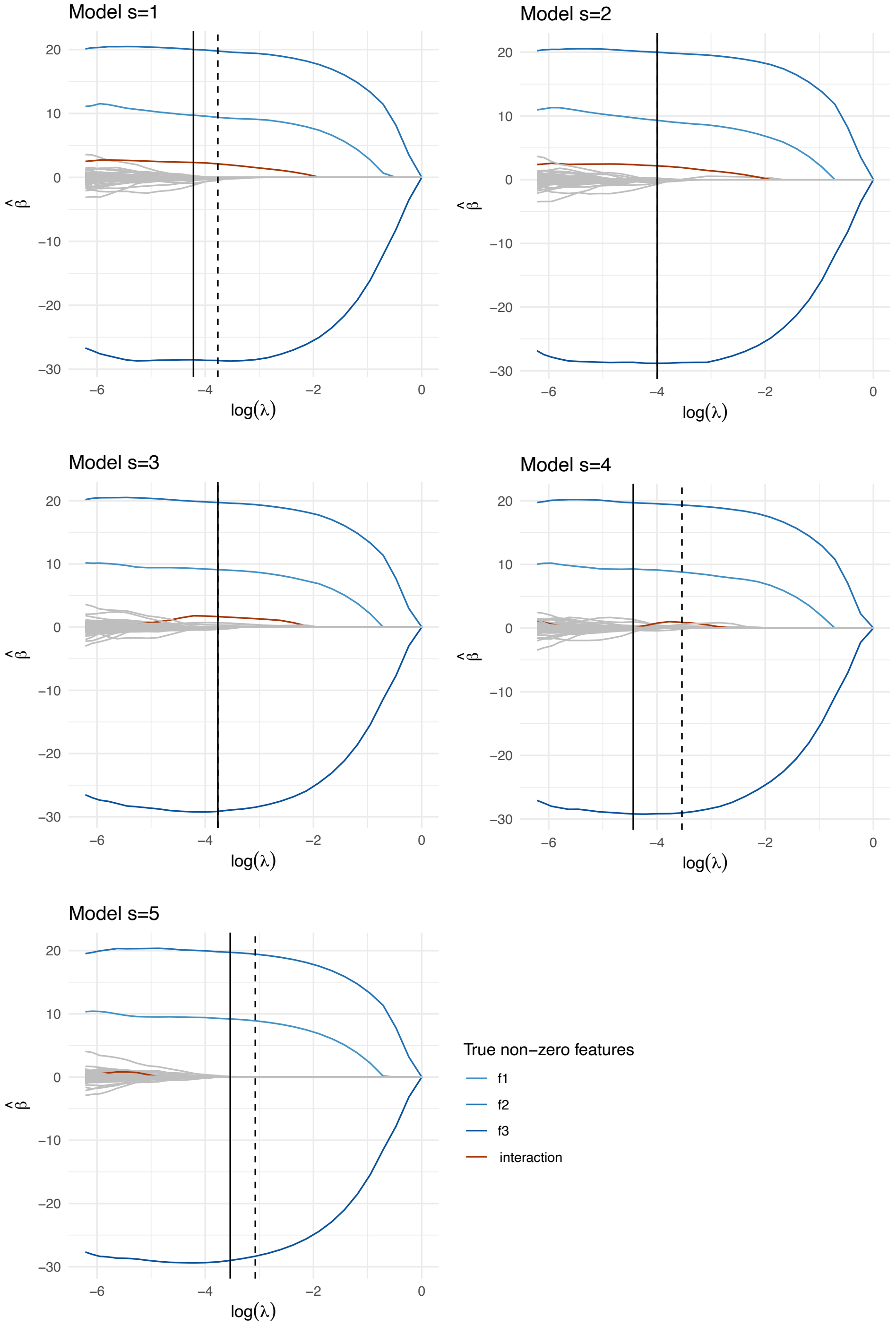


**Fig S4.** Solution path of the interaction model (sparse qlc) for the $S = 5$ semi-synthetic simulation setups for varying feature sparsity levels.

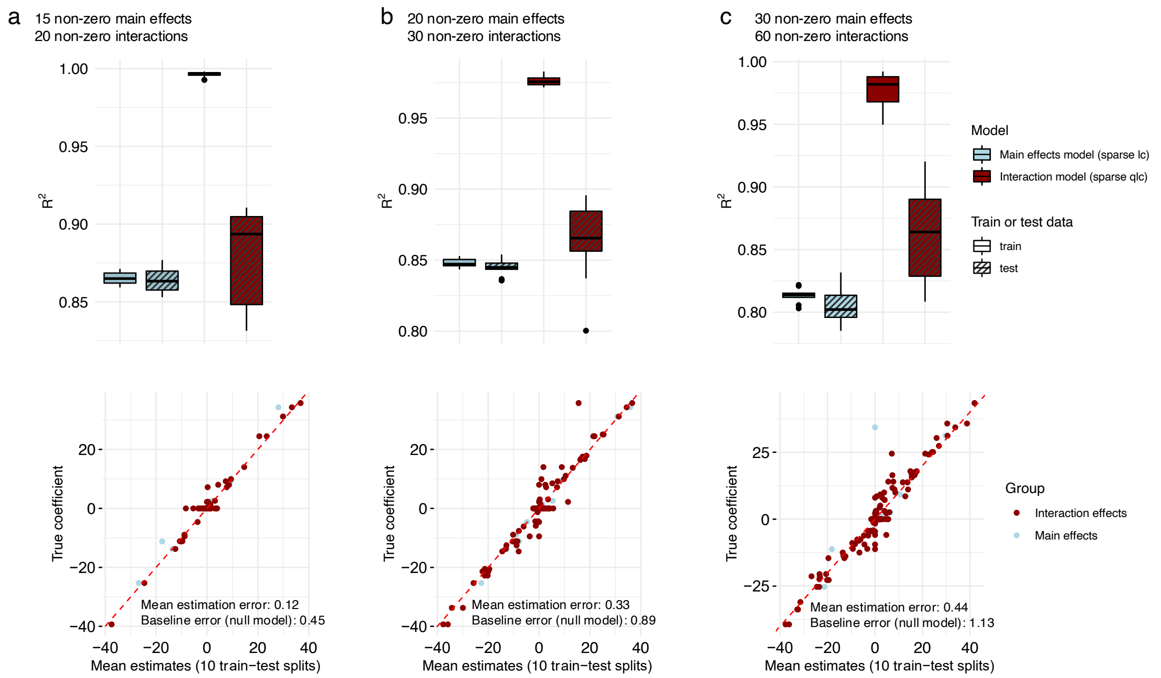


**Fig S5.** Semi-synthetic data simulations according to the sparse quadratic log-contrast model based on the American Gut Project data at the family level from $A_{\text{AGP}}$, comprising $n = 6266$ samples and the $p = 50$ most prevalent families, leading to a compositional count matrix $A_{\text{AGP}} \in \mathbb{R}_{+}^{6266 \times 50}$. We fix the intercept term at $\beta_0^* = 10$ and vary the number of non-zero main and interaction effects. The non-zero entries are sampled from a normal distribution $\beta^* \sim \mathcal{N}(0, \sigma)$ and $\Theta^* \sim \mathcal{N}(0, \sigma)$. Panels (a, b, and c) show the distributions of the estimated coefficients across 10 train-test splits, highlighting the effect of varying the number of 15 to 30 non-zero main effects (a: 15, b: 20, c: 30) and 20 to 60 interaction coefficients (a: 20, b: 30, c: 60).
